# Wetting and complex remodeling of membranes by biomolecular condensates

**DOI:** 10.1101/2022.06.03.494704

**Authors:** Agustín Mangiarotti, Nannan Chen, Ziliang Zhao, Reinhard Lipowsky, Rumiana Dimova

## Abstract

Cells compartmentalize their components in liquid-like condensates, which can be reconstituted *in vitro*. Although these condensates interact with membrane-bound organelles, the potential of membrane remodeling and the underlying mechanisms are not well understood. Here, we demonstrate that interactions between protein condensates (including hollow ones) and membranes can lead to remarkable morphological transformations and describe these with theory. Modulation of solution salinity or membrane composition drives the condensate-membrane system through two wetting transitions, from dewetting, through a broad regime of partial wetting, to complete wetting. A new phenomenon, namely fingering or ruffling of the condensate-membrane interface is observed when sufficient membrane area is available, producing intricately curved structures. The observed morphologies are governed by the interplay of adhesion, membrane elasticity, and interfacial tension. Our results highlight the relevance of wetting in cell biology, and pave the way for the design of synthetic membrane-droplet based biomaterials and compartments with tunable properties.

## Introduction

The last decade of research has provided ample evidence that, in addition to membrane-bound organelles, cells compartmentalize their interior by membraneless organelles also referred to as macromolecular condensates or coacervates, which behave as liquid droplets. Examples include nucleoli, Cajal bodies, P-bodies, and stress granules, all of which represent liquid protein/RNA-rich droplets within the cell^1, 2, 3^. They arise from the condensation of cellular material through liquid-liquid phase separation and can be reconstituted *in vitro*^4, 5, 6^. This discovery has expanded the search for the different functions of liquid droplets in animal, fungal, and plant systems, which include compartmentalization, sorting of macromolecules, tuning of enzymatic reactions, preservation of cellular fitness, immune response, and temperature sensing^1, 3, 7^. In addition, an aberrant protein condensation has been proposed as an intermediate step in neurodegenerative diseases such as Parkinson and Alzheimer^3, 8^. Thus, the research on membrane-less organelles has become a very active and strongly interdisciplinary research field^7^.

One interesting and hardly explored aspect of these liquid droplets is that, although termed “membraneless”, they can come into contact and wet membranous compartments^9, 10^. In the last years, it has been found that membrane-droplet interactions are involved in biological key processes such as signal transduction in T cells^11^, the assembly of virus capsid proteins^12^, ribonucleoprotein granules biogenesis and fission in the endoplasmic reticulum^13, 14^, the development of tight junctions^15^, and the assembly of endocytic vesicles^16^. However, a detailed understanding of the underlying physicochemical mechanisms and a systematic characterization of the membrane-droplet interactions is still missing because these interactions are difficult to assess *in vivo*, to some extent reflecting the small size of the condensates^17^.

So far, membrane-droplet interactions have been studied in the context of only a few *in-vitro* systems. When aqueous two-phase systems (ATPS) consisting of polymer solutions^18, 19, 20, 21, 22^ are encapsulated or in contact with giant unilamellar vesicles (GUVs)^23^, several biologically relevant processes were demonstrated to occur, namely, outward or inward budding mimicking exo- and endocytosis processes, formation of membrane nanotubes and fission of vesicle compartments^22, 24^. The transitions were triggered either thermally or via osmotically-imposed concentration changes. Another approach consisted of inducing coacervation in the GUV interior by externally changing the pH^25^. The uptake of coacervate droplets by vesicles via endocytosis was demonstrated to be modulated by electrostatic droplet-membrane interactions^26^. Furthermore, membrane tubulation was observed in GUVs decorated with anchored proteins that underwent phase separation^27^.

While several studies have already addressed the effect of membrane composition and phase state on the interaction with condensates^27, 28^, as well as the coupling between lipid domains and condensates^29, 30^, we focused on understanding the mechanism of membrane wetting and remodeling by condensates. Here, we provide a systematic analysis of the membrane remodeling and wetting behavior of GUVs exposed to water-soluble proteins that phase separate into a protein-rich and a protein-poor phase. As a model protein, we employed glycinin, which is one of the most abundant storage proteins in the soybean. Glycinin undergoes liquid-liquid phase separation in the presence of sodium chloride^6, 31^, making it a convenient model system to study the membrane-droplet interaction. When combining glycinin condensates with GUVs we observed condensate-membrane adhesion, partial and complete wetting, condensate-induced budding, and a remarkable new feature of this interaction: the complex remodeling of the membrane-droplet interface producing ruffled membrane-condensate structures with fingers. Furthermore, shifts within the phase-coexistence region of the glycinin phase diagram, generate hollow condensates, which can also spread on the membranes, thus expanding possibilities for cell-mimetic compartmentalization. These diverse phenomena and morphologies are summarized in Figure 1. We demonstrate that our findings do not only apply to glycinin condensates, but also to peptide coacervates and PEG/dextran ATPS. Our study has important implications for the droplet-induced morphogenesis of ubiquitous membrane-bound organelles, and, in addition, for material science and synthetic biology in the development of smart soft materials with compartmentation and morphology modulated by wetting.

**Figure 1:**
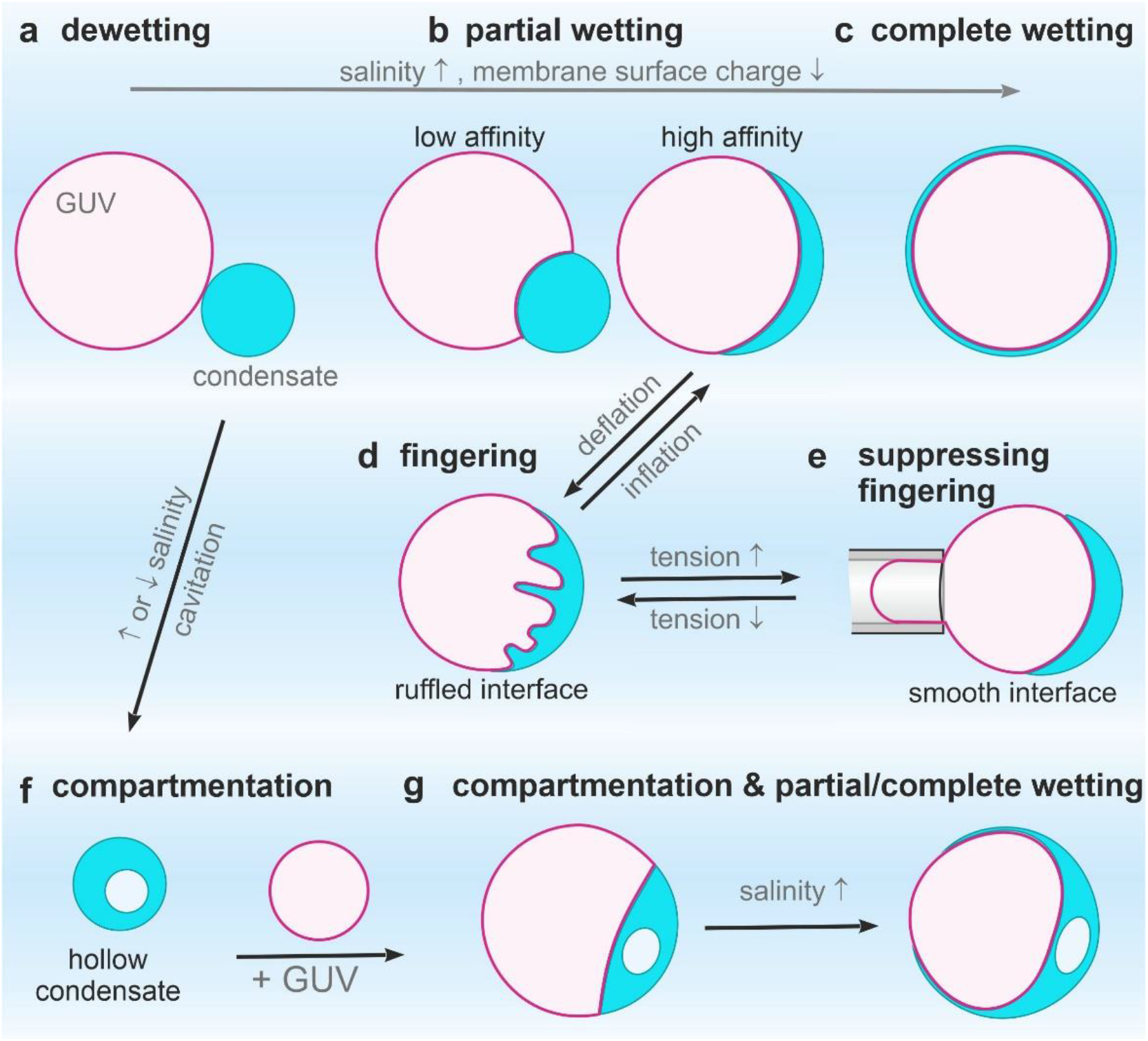
Membranes in contact with biomolecular condensates can undergo two wetting transitions from dewetting (a, f) through partial wetting (b, g-left) to complete wetting (c, g-right) modulated by ionic strength and membrane composition. Complex morphological transformations exhibited by interface ruffling and fingering (d) can be observed when excess membrane area is available, or suppressed when this area is retracted upon tension increase (e). Salinity changes can result in the formation of hollow condensates (f), which can also exhibit partial and complete wetting (g) offering additional means of compartmentation in cells.

## Results

### Wetting transitions at membranes in contact with biomolecular condensates

Liquid droplets at surfaces adopt different shapes depending on the strength of interaction: they may remain spherical (no wetting), or slightly spread on the surface adopting the morphology of a truncated sphere (partial wetting), or completely spread, wetting the whole surface (complete wetting). By changing control parameters such as surface or droplet composition, temperature or interaction strength, the system can undergo a transition from dewetting to partial wetting and even complete wetting^32, 33^. Similarly, when biomolecular condensates get in contact with membranes, they can undergo wetting transitions depending on the interaction strength^19^. However, contrary to solid substrates, membranes can deform because of their relatively low bending rigidity. Membrane wetting transitions were first described for vesicles in contact with ATPS composed of a mixture of poly(ethylene glycol) (PEG) and dextran^19^. In these systems, several morphological transformations have been produced by modulating the wetting of the membrane by the polymers ^21, 34^. Figure 2a shows an example of a partial wetting morphology for a vesicle in an ATPS system. An out-wetting morphology can be observed where the condensate droplet (green) is from the dextran-rich phase; similar examples for glycinin condensates in contact with membranes are shown in Fig. 2e. To quantify the membrane-condensate interactions, we first provide a theoretical description of the system (see Methods for details): in general, partial wetting morphologies involve three different surface segments, which meet along the contact line (Fig. 2b), as well as three contact angles and surface tensions (Fig. 2c)^34, 35, 36, 37^. The contact angles and the mechanical tensions 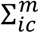 and 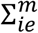 of the two membrane segments depend on the lateral stress Σ within the membrane (Methods, Eq. 8) and thus on the size and shape of the vesicle. In contrast, both the interfacial tension Σ_*ce*_ and the affinity contrast *W* (Methods, Eq. 9), which represents the difference between the adhesion free energies per unit area of condensate and external buffer, are material parameters, which do not depend on the size and shape of the chosen condensate-vesicle couple. Furthermore, the ratio of the affinity contrast *W* to the interfacial tension Σ_*ce*_ is directly related to the three contact angles and, thus, can be obtained by measuring these angles from the microscopy images (Methods, Eq. 11, see also Fig. 3b). Using this geometric and membrane-elastic description, we systematically studied the interaction of glycinin condensates^6^ with GUVs made of zwitterionic phosphatidylcholine and GUVs enclosing PEG-dextran ATPS. Upon interaction of the condensates with the membranes, the system undergoes two wetting transitions (see Figs. 2d-e, S1a,c,d for data on glycinin and varied membrane composition; analogous trend is observed upon raising ATPS polymer concentrations, see Fig. S1b). At low salt content, the glycinin droplets do not interact with the vesicles (dewetted state, Φ = 1). Raising the salinity leads to increased attractive interaction between the droplets and the membrane (partial wetting, −1 < Φ < 1) until complete wetting (Φ = −1), i.e., spreading of the condensates over the whole membrane surface, which is reached when the salinity exceeds 180 mM NaCl (Figs. 2d-e, S1a,d). The observed behavior plausibly results from charge screening of the protein droplets (glycinin is negatively charged at the working pH=7), allowing for the interaction with the membrane. The experimental data show that glycinin condensates interacting with GUVs exhibit three different wetting regimes separated by two wetting transitions at a constant temperature. Even though wetting transitions have been extensively studied in the past, see e.g. ^33, 38, 39^, to the best of our knowledge, this behavior has not been previously reported ascribing remarkable features to this condensate-membrane system.

**Figure 2:**
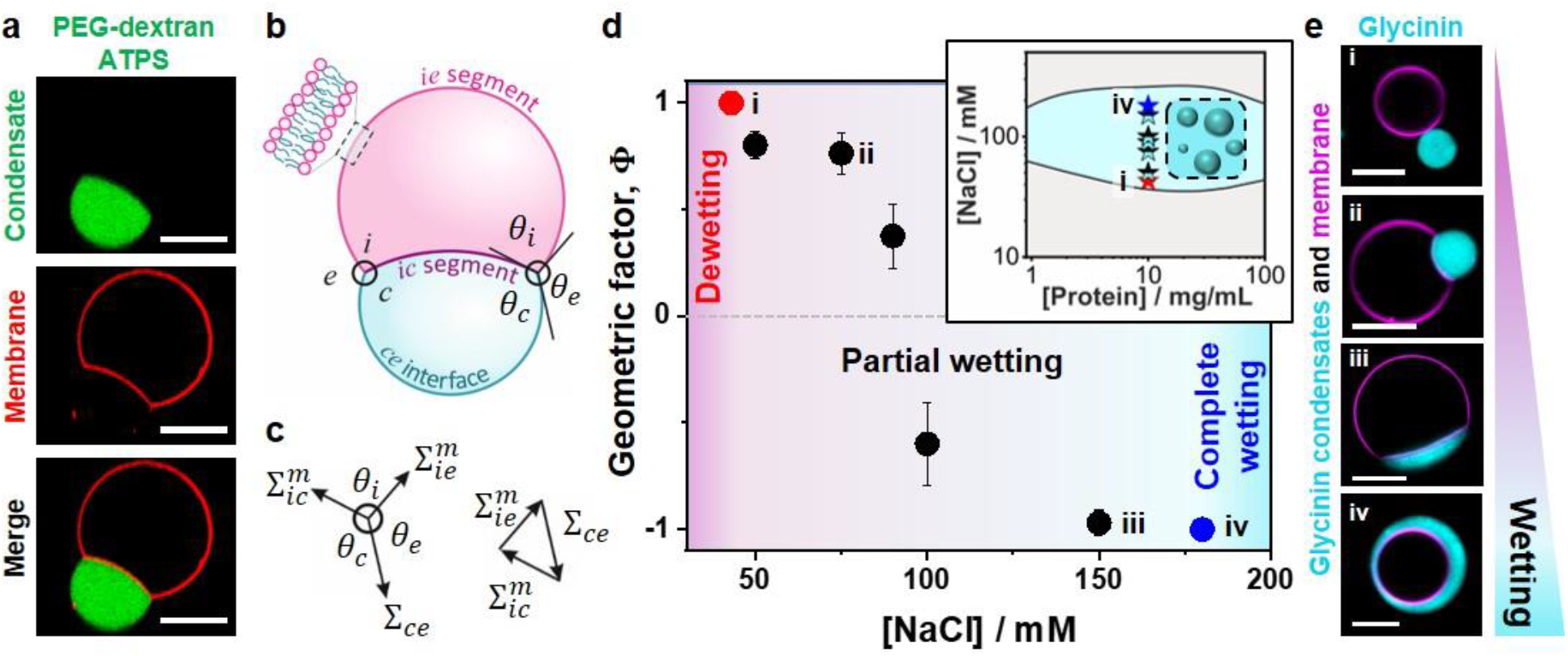
Membrane wetting by PEG/dextran and glycinin condensates. **(a)** Confocal microscopy images showing partial out-wetting of a vesicle (red) by a dextran-rich droplet (green) in phase-separated PEG/dextran aqueous two-phase systems; the vesicle adopts a moon-like shape. **(b)** Partial wetting leads to the formation of a contact line between the *ce* interface (blue line) and the membrane. The contact line partitions the membrane into the *ie* (magenta) and *ic* (purple) segments, with the contact angles *θ*_*c*_ + *θ*_*i*_ + *θ*_*e*_ = 360°. **(c)** Force balance between the droplet interfacial tension Σ_*ce*_ and the mechanical tensions 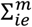 and 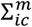 within the two membrane segments. These three tensions form the sides of a triangle (which implies Eqs. 9 and 10 in Methods)^40^. **(d)** Experimental data for the geometric factor Φ = (sin 𝜃_*e*_ – sin 𝜃_*c*_)/sin 𝜃_i_ (Eqs. 8-11 in Methods) as a function of NaCl concentration. The largest possible value Φ = +1 (red circle) corresponds to the transition from dewetting to partial wetting at [NaCl]≈43 mM, the smallest possible value Φ = −1 (blue circle) reflects the transition from partial wetting to complete wetting at [NaCl]≈180 mM. All data: mean ± SD, n=10 per composition. Data for the geometric factor of the PEG/Dextran system can be found in Fig. S1b. The inset shows the phase diagram of glycinin in salt solutions with the region of phase coexistence shown in cyan. The locations of the measured geometric factor data points are indicated with stars. **(e)** Examples of confocal microscopy images of different wetting morphologies for the data points indicated in (d); for more images see Fig. S1a. All scale bars: 10 µm.

**Figure 3:**
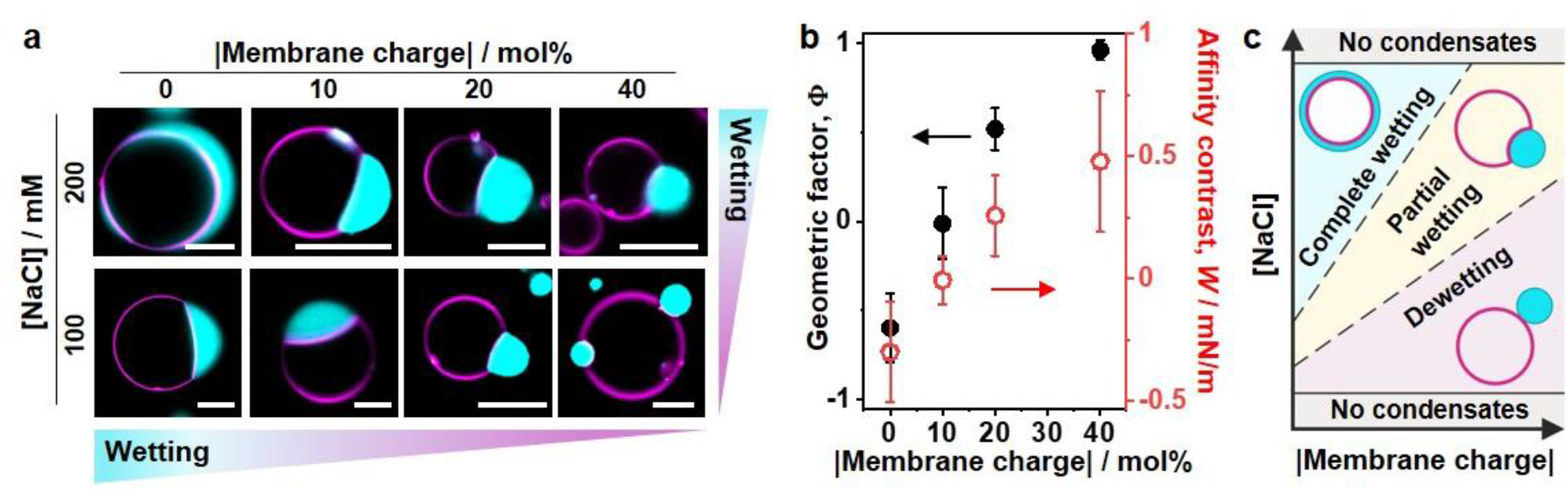
Tuning wetting by membrane charge and salinity. **(a)** Confocal microscopy images of the vesicle-droplet system showing characteristic wetting morphologies as a function of salinity and membrane composition (in terms of absolute value of the membrane charge), see Fig. S2 for more data. Scale bars: 10 µm. **(b)** Geometric factor Φ (black axis) and affinity contrast *W* (red axis) vs absolute membrane charge (% mol) for giant vesicles in contact with glycinin condensates at 100 mM NaCl (data for 10mol% DOPS and 10mol% DOTAP are combined, see Fig. S2). The affinity contrast is given in units of interfacial tension (*W* = Φ Σ_*ce*_, see Methods) and the errors are estimated with error propagation analysis. All data: mean ± SD, n=10 per composition. **(c)** Schematic morphology diagram of the droplet-vesicle interactions as a function of salinity and membrane charge.

Noteworthy, the wetting transitions can be easily tuned not only by changes in salinity, but also by modifying the membrane composition (Figs. 3, S2). When increasing the absolute membrane charge by including negatively (DOPS) or positively charged lipids (DOTAP), the system moves towards dewetting (Fig 3a,b). This behavior is counterintuitive, because one might have expected that increasing the positive charge in the membrane would lead to a stronger interaction with the negatively charged protein; but the opposite is observed. In both cases, for negatively and positively charged membranes wetting decreases as compared to the affinity for neutral membranes, highlighting the complexity of the glycinin condensates and suggesting that their interaction with the membrane is not only electrostatic in nature but presumably also hydrophobic. In fact, previous reports have described favorable hydrophobic interaction between glycinin hydrophobic residues and phosphatidylcholines or lecithin^41, 42^. It is important to note that glycinin is a relatively large molecule (a hexamer, roughly 5 nm in size) presenting acidic and basic residues that undergo conformational changes upon phase separation^6, 43^. Altogether, this suggests the involvement of hydrophobic interactions of glycinin with membranes, which are enhanced at higher fractions of neutral lipids. With increasing salinity in these systems, membrane charge is screened (Figs. 3a and S2), favoring wetting. In this manner, membrane composition and salinity are simple parameters that allow for fine-tuning of the wetting morphologies and stability of condensates (Fig. 3c).

Inspection of the partial wetting morphologies in Fig. 2 and 3 shows that the visible membrane segments *ie* and *ic* are separated by an apparent kink in the membrane shape (see for example the cusp of the moon-like vesicle seen in the membrane channel in Fig. 2a). The fine structure of this kink corresponds to a highly curved membrane segment, which cannot be resolved in the confocal microscope (but visualized with STED microscopy^44^). It arises from the mechanical balance between the capillary forces, generated by the interfacial tension Σ_*ce*_ of the condensate-buffer interface, and the membrane bending moment which is proportional to its bending rigidity *κ*. This mechanical balance implies that the contact line segment acquires a curvature radius of the order of 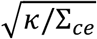. For typical values of the bending rigidity *k ⋍* 10^−19^ J and the interfacial tension Σ_*ce*_ *⋍*0.5 mN/m (measured here, see Methods), we obtain the curvature radius 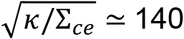, which is indeed below the optical resolution limit and thus appears as a kink in the confocal images. The length scale 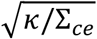 enters the normal (or perpendicular) component of the force balance between the three surface tensions at the contact line as shown by minimization of the combined curvature and adhesion energy of the vesicle-droplet couple^35^. In the context of liquid droplets at the surfaces of solid materials, the competition between the surface tension Σ_int_ of the liquid-vapor interface and the Young’s modulus ϒ of the solid material leads to the elastocapillary length Σ_*int*_/ ^ϒ^ ^45, 46^. Note that the elastocapillary length grows with increasing surface tension whereas the curvature Radius 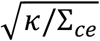 decreases with increasing interfacial tension.

### Droplet spreading and dynamics

In all cases, membrane wetting by the protein condensates was characterized by slow dynamics in the minutes-to-hour range (Fig. 4a, Movie S1). Once in contact, the angles characterizing the condensate-membrane system required 20-30 min to reach equilibrium (Fig. 4b). This is most likely related to the high molecular weight of glycinin (360 kDa) that translates in the high viscosity of glycinin condensates ∼4.8 kPa s (see Methods), which is roughly three orders of magnitude higher than that of condensates formed by other proteins like FUS (0.7 Pa.s.) or PGL-1 (1 Pa.s)^47^. The data in Figs. 2 and 3 as well as the results described in the following correspond always to the equilibrated final wetting morphology.

**Figure 4:**
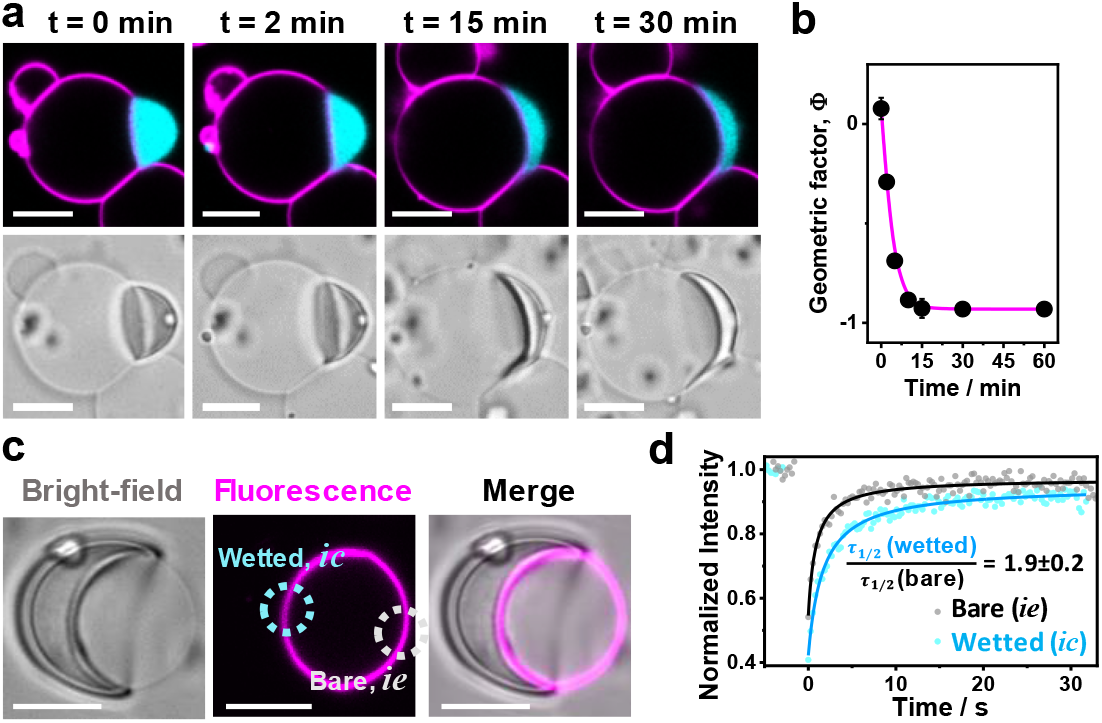
Wetting and membrane dynamics: **(a)** After the glycinin condensate touches the membrane (t=0 min), the contact angles change slowly until reaching the final morphology, which remains stable for hours; confocal (top) and bright-field (bottom) images of the same vesicle-condensate pair. **(b)** Measured geometric factor Φ for the membrane-condensate system shown in (a); DOPC membrane, 100 mM NaCl. **(c)** Fluorescence recovery after photo-bleaching (FRAP) experiments show decreased fluidity in the membrane segment that is wetted by the condensate (*ic* segment in Fig. 2b) compared to the bare one (condensate-free, *ie*) on the same vesicle. For these experiments, only the membrane was fluorescently labeled to avoid interferences from condensate fluorescence. The dotted circles shown in cyan/gray indicate the bleached regions in the *ic*/*ie* membrane segments respectively. **(d)** FRAP intensity curves yield halftimes of recovery *τ*_1/2_ which show that condensate wetting slows lipid diffusion by a factor of about 2 (n = 5); see Methods. Scale bars: 10 μm.

Compared to bare (condensate-free) membranes (segment *ie* in Fig. 2b), vesicle segments in contact with condensates (segment *ic*) are characterized by lower fluidity as a result of the protein-lipid interaction, with the immobile fraction in such membranes being 16±5% measured by FRAP (Fig. 4c,d). This indicates that the condensates not only induce membrane morphological transformations, but also impose dynamic constraints onto the constituting lipid species.

### Hollow condensates and droplet-vesicle bridging provide means for compartmentation

Similarly to the behavior observed for the PEG-dextran ATPS system in contact with membranes, glycinin condensate-vesicle interactions can lead to complete droplet engulfment (as long as sufficient membrane area is available), a vesicle can bridge several condensates or a condensate can enclose or bridge several GUVs. The necessary condition for engulfment is (partial or complete) wetting and excess vesicle area (compared to the area of a sphere enclosing the same volume) sufficient to wrap the condensate. Thus, wetting can give rise to complex condensate-vesicle architectures of multiple compartments (Figs. 5, S3), demonstrating the remodeling role of capillary forces^10, 37^. As recently reported for coacervates^26^, the engulfment of condensates by the vesicle membrane can separate the individual droplets from each other and prevent their coalescence. In addition, this process also changes the area of the interface between the condensate and the liquid bulk phase, thereby regulating the diffusive exchange of molecules between the droplets and the bulk phase. Remarkably, hollow condensates, obtained by inducing phase separation within the protein-rich droplets by shifting the salinity within the coexistence region of the phase diagram^6^ (see Fig. 5b,c and Methods), can reshape the membranes in a similar way (Fig. 5d), thereby providing additional means of compartmentation. Considering the abundance of cellular membranes in the cell interior, we speculate that bridging, separation, and further compartmentation interactions mediated by membranes are highly relevant.

**Figure 5:**
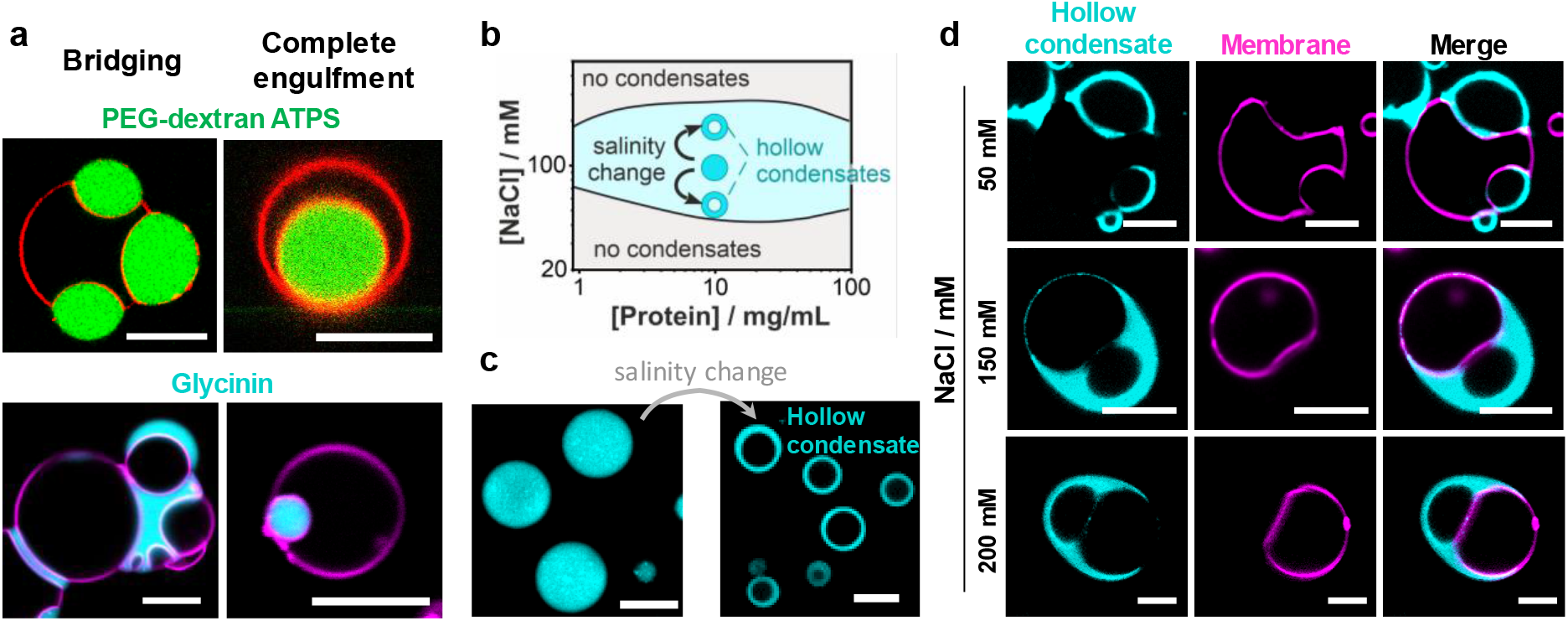
Complex architectures and compartmentation generated by condensate-membrane interactions: **(a)** Bridging and complete engulfment in condensate-membrane systems, where the condensate droplets are either dextran-rich (green) in PEG-dextran ATPS (upper row) or glycinin condensates (lower row). Either the vesicle membrane can bridge several droplets (top, left) or a condensate can bridge several vesicles (bottom left) creating complex architectures with several compartments. The engulfment of the condensate (right images) is energetically favored by both the increased contact area with the membrane and the reduced surface area of the *ce* interface (see Fig. 2b). **(b-d)**. Hollow condensates formation and membrane wetting. **(b)** Schematic illustration of the pathways in the glycinin phase diagram used to generate hollow condensates starting from droplets prepared in 100 mM NaCl and exposed to salinity shift to higher or lower concentration of NaCl, which triggers phase separation within the droplets. **(c)** Images of the condensates before and after the salinity change; here, from 100 mM NaCl (left) to 50 mM NaCl (right), showing the formation of hollow condensates by such a salinity downward shift. **(d)** Hollow condensates (cyan) in contact with soy-PC GUVs (magenta) at the indicated NaCl concentrations allow for additional system compartmentation (see also Fig. S3). All scale bars are 10 μm.

### Ruffling and fingering of the condensate-membrane interface

Probably the most spectacular response to wetting in the glycinin-membrane system is observed on vesicles with excess area: the membrane-condensate interface (*ic* segment in Fig. 2b) undergoes ruffling which can proceed to excessive curving of the membrane/droplet interface (Fig. 6). This process of interfacial ruffling is reminiscent of viscous fingering observed when less viscous liquid displaces a more viscous one in porous media^48^, although here a membrane separates the fluids, and pressure and porous confinement are absent. An even more relevant analogy is provided by the structural resemblance to elements of the endoplasmic reticulum, despite the missing network-like character (which could be created by subsequent fusion of the protrusions as generated e.g. by fusion proteins) and slightly higher curvature (imposed by adsorbing or scaffolding proteins such as reticulons and atlastins).

**Figure 6:**
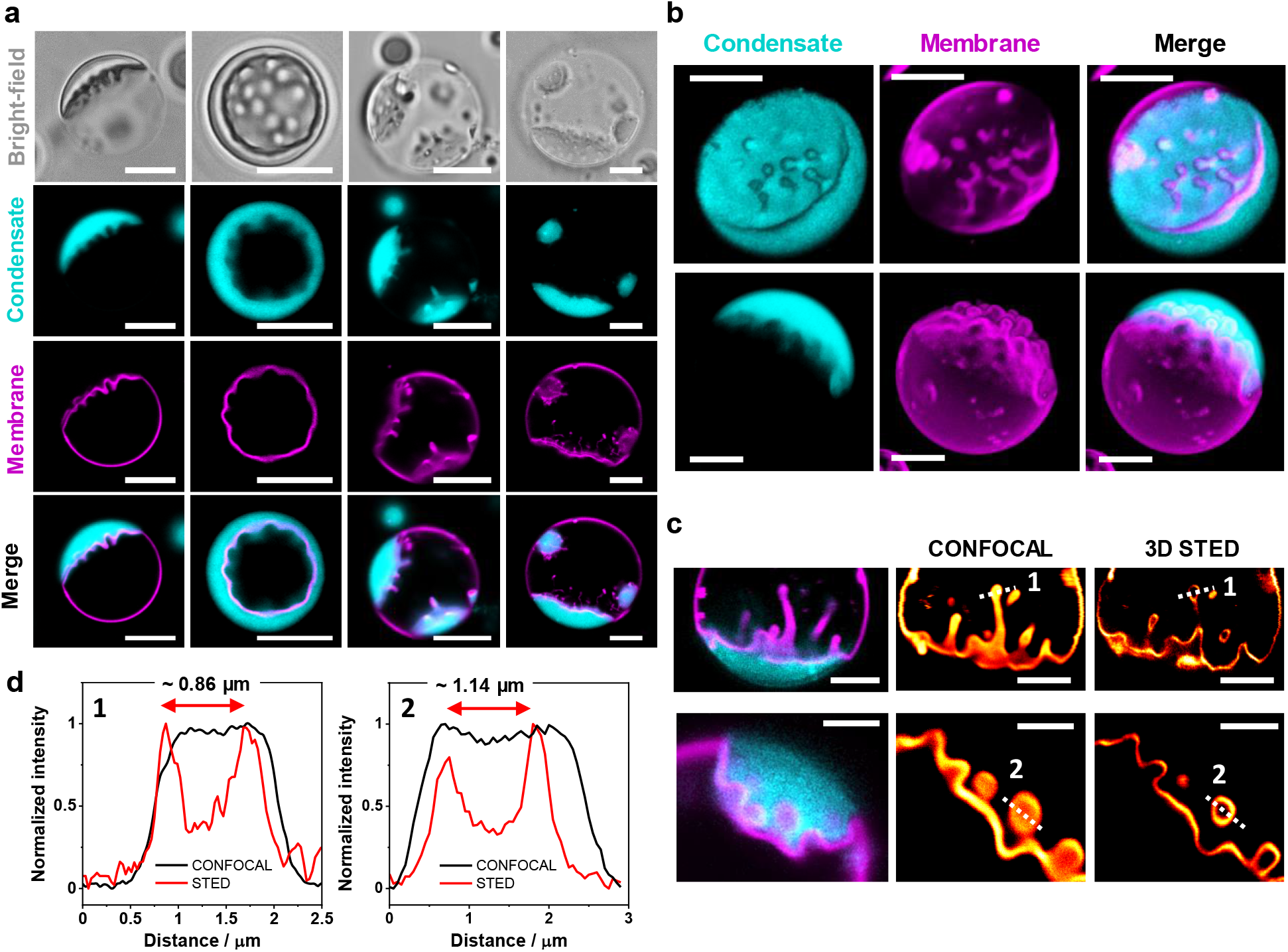
Ruffling and fingering of the membrane-condensate interface leads to complex morphologies: **(a)** Examples of ruffling morphologies of the condensate-membrane interface seen in bright field and confocal cross-sections. **(b)** 3D reconstructions of GUVs in contact with glycinin condensates showing the different ruffled morphologies (see also Fig. S4). **(c)** 3D STED imaging resolves characteristic features of the fingered regions in the submicrometer range (see also Fig. S5). Intensity line profiles along the indicated dashed lines in the micrographs in (c) allow for an estimation of the finger thickness (left corresponds to the first image and right to the second one as indicated by the numbers in panel c). All scale bars: 5 µm.

The complex membrane shapes include large protrusions and invaginations in the form of finger-like structures and thin tubes (Fig. 6a-c, Movie S2). It can be seen that the ruffling/fingering involves the mutual scaffolding of membrane and condensates, since both become curved (Figs. 6a,b, S4a), and the behavior is observed also for more complex membrane compositions (Fig. S4b). The phenomenon is different from membrane tubulation triggered by spontaneous curvature generation as observed in membrane wetting by ATPS systems, where tubes were observed to protrude always towards the PEG-rich phase (see e.g.^21, 44, 49^). Similarly, tubulation via compressive stress induced by protein phase separation at the membrane surface^27^ can also be excluded because of the lack of directionality in the protrusions observed here. We observe that the interface can develop fingers to either side, presumably depending on the volume of the droplet or vesicle provided for the fingerling. Using 3D STED microscopy, we resolved characteristic features and dimensions of the reticulated (elastically deformed) regions which lie in the submicron-micron scale (Figs. 6c-d, S5).

The process of membrane ruffling and fingering is slow. It initiates while the contact angle is stabilizing (see Fig. 7a,b, Movie S3) and completes within 40-60 min. Note that vesicles exhibiting ruffled or fingered membranes were not considered when assessing the geometric factor Φ in Figs. 2-4, because in such systems the contact angles are ill-defined. Once established, the ruffled structure does not fluctuate (Movie S4) and remains stable for hours. As shown in Fig. 7a (and Movie S3), the contact area between the membrane and the condensate increases with the degree of ruffling.^**min t = 10 min t = 20**^ The intricate and striking membrane morphologies arise from the gain of adhesion energy between droplet and membrane when excess area is pulled out of the *ie* segment and added to the *ic* segment, see Methods. Subsequently, the tension of this segment becomes negligible 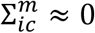, which implies, via the force balance triangle (Fig. 2c) that 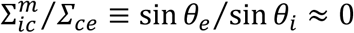 and that the angle *θ*_*e*_ should approach 180°. This is indeed consistent with our experimental observations that show that the *ic* membrane segment becomes parallel to the *ce* interface along the contact line, i.e. the vesicle-condensate pair rounds up overall (see e.g. Figs. 6a, 7c). Furthermore, this tension relation together with Eqs. (10) and (11) imply negative values for the geometric factor Φ, and indeed, interface ruffling was observed only for [NaCl] ≥ 100 mM where Φ < 0, i.e. for negative affinity contrast. Whether or not ruffling/fingering is associated with an increase in the bending energy depends on the spontaneous curvature of the *ic* segment. The adhesion of the glycinin-rich droplet implies adsorption of the protein to the membrane, for which one would expect the membrane to bulge towards the droplet as is the case for Φ < 0. In this case, adhesion and bending act in a synergistic manner. At their liquid-liquid interfaces, condensates exhibit some degree of molecular organization as shown previously by birefringence^6, 50^. We cannot exclude that this surface organization is perturbed by the interaction of the membrane with the proteins at the droplet interface. However, the lipid mobility in the reticulated membrane segment remains similar to that in smooth membrane-condensate segments (compare Figs. 7c,d and 4c,d). We also confirmed that the membrane integrity is preserved during wetting and ruffling (Fig. S6). Furthermore, considering that solution asymmetry (in terms of salinity) across the membrane, as is the case for the GUVs examined here, can drive tubulation^51^, we explored whether the interfacial ruffling will be preserved when exposing the membrane to symmetric salt conditions. We confirmed that ruffling is independent of asymmetric buffer conditions (Fig. S7).

**Figure 7:**
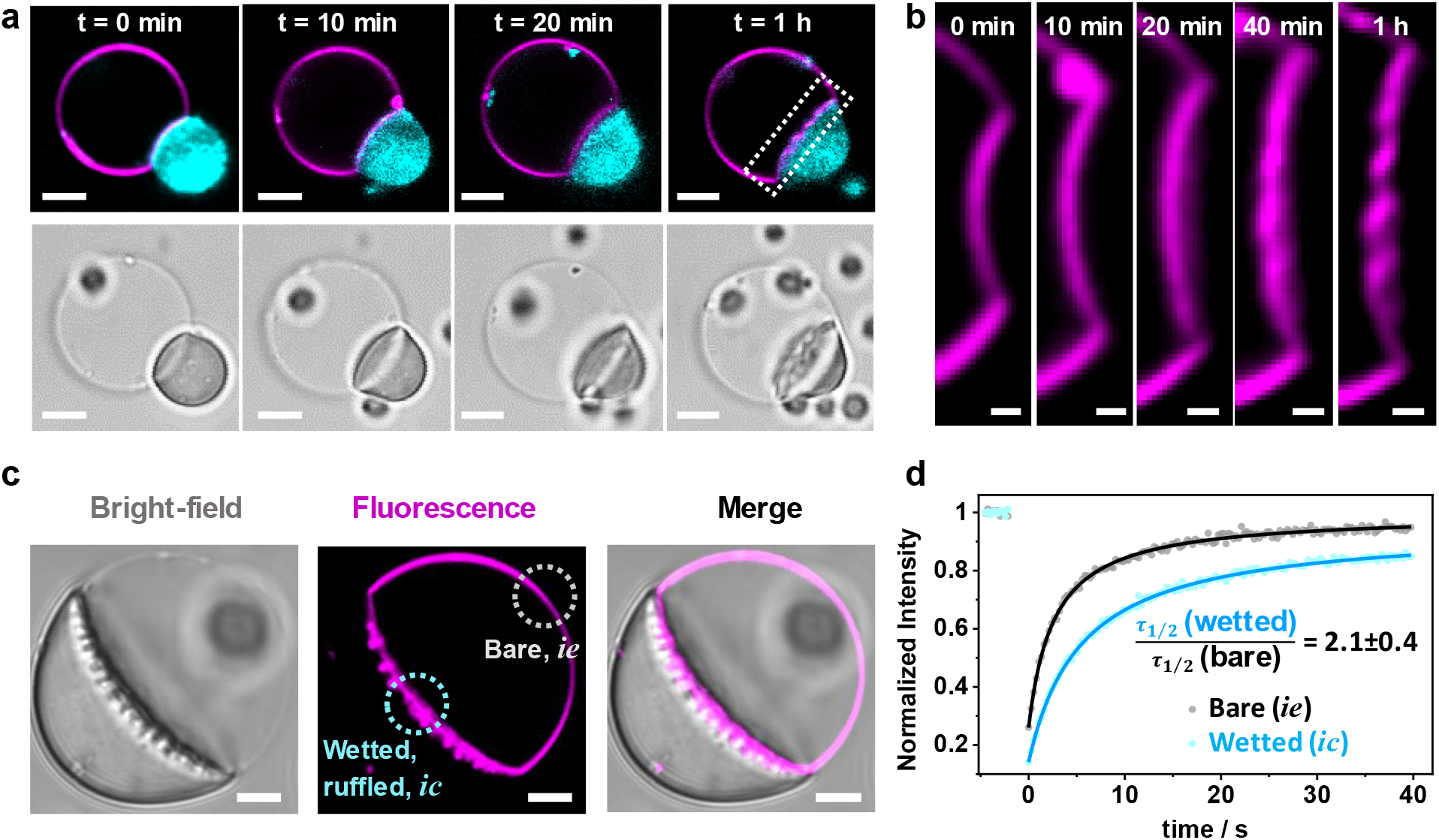
Dynamics of membrane ruffling driven by glycinin condensates: **(a)** Mutual membrane-condensate molding and generation of curved structures at the membrane-droplet interface during wetting proceeds slowly (see also Movie S3). **(b)** Zoom of the region indicated with a dotted line in (a). **(c)** Images in bright field and confocal cross section showing the two membrane segments examined by FRAP, the bare one (*ie* segment in Fig. 2b) and the wetted, reticulated one (*ic*). Only the membrane was labeled and it was bleached in the indicated regions. **(d)** Recovery curves for the reticulated region (cyan) and the bare condensate-free membrane (grey). From the fitting, the halftimes of recovery τ_1/2_ are obtained (n=5, see Methods). Lipid diffusion in the membrane in contact with the condensate is twice as slow than in the condensate-free segment similarly to measurements on smooth interfaces where the slowdown is the same (compare to Fig. 4c,d). These results indicate that membrane ruffling by condensates does not alter lipid diffusion compared to that in wetted membranes where ruffling is absent. Scale bars in (a) and (c): 5 µm, in (b): 1 µm.

As indicated above, the remarkable interface ruffling and finger-like structures are only present in vesicles with available excess area. We explored the vesicle response to osmotic deflation and inflation (see details in Methods). When vesicles are deflated by increasing the external osmolarity they gain excess area (compared to the area of a sphere with the equivalent volume) and either exhibit visible fluctuations (quasi-spherical or floppy vesicles) or deform into non-spherical shapes, where the excess area could be stored in the form of buds or nanotubes stabilized by spontaneous curvature^40, 52^. In contrast, when decreasing the external osmolarity, vesicles become tense and spherical, thus suppressing fluctuations. Interfacial ruffling was increasingly more pronounced upon stronger osmotic deflation (Fig. 8a). The effect of membrane excess area becomes even more evident when the membrane tension is increased via micropipette aspiration (Methods), which suppresses ruffling leading to smooth vesicle-droplet interfaces (Fig. 8b, Movie S5). Reversely, decreasing the tension (suction pressure) restores ruffling. The tensions (∼1 mN/m) at which the interface smoothens are much higher than those needed to retract tubes formed in ATPS-vesicle systems (∼20-100 μN/m)^49^. This difference could be presumably attributed to the much higher interfacial tension of the protein condensates (∼0.5 mN/m, Methods) compared to that in the ATPS system (∼0.01 mN/m)^53^.

**Figure 8:**
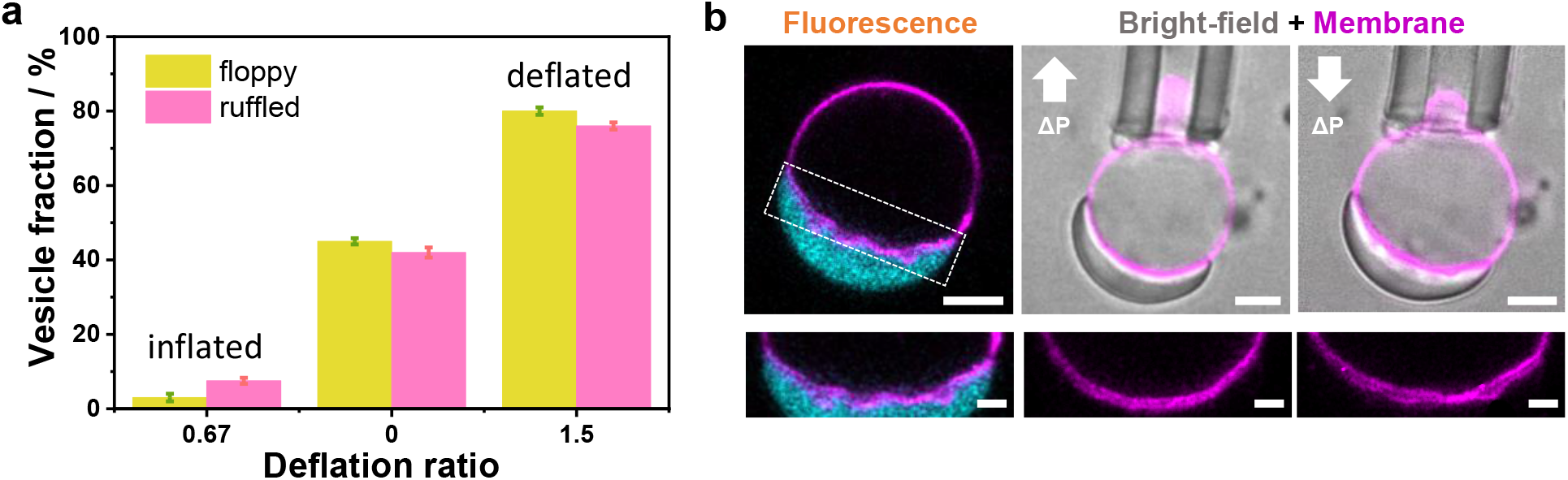
Ruffling of the membrane-condensate interface is enhanced for vesicles with more excess area and can be tuned by tension. **(a)** The percentage of vesicles in which ruffling is observed increases with increasing the membrane excess area as shown by the fraction of floppy vesicles. Deflation ratio in the graph is the ratio of the externally applied osmolarity to the initial vesicle osmolarity (n=60 for each condition and the values shown are mean ±SD of three independent experiments). **(b)** Increasing membrane tension via micropipette aspiration results in the suppression of ruffles and smoothing out of the condensate-membrane interface. The tension threshold needed to suppress the ruffles is (1.0±0.2) mN/m (n=5). When releasing the membrane area by decreasing the tension, ruffles reappeared within a minute (upper middle and right panels, up- and down-arrows signify increasing and decreasing suction pressure). Scale bars are 5 µm. A zoom of the membrane region highlighted with white dashed line is shown in the lower panels. Scale bars are 2 µm.

The above results on membrane remodeling and wetting transitions were obtained using glycinin droplets, but biomolecular condensates can exhibit a wide range of surface charge, interfacial tension and viscosity^47^. It is important to note that the wetting state can be expected to depend not only on the material properties of the condensate, but also on the specific interactions between particular lipids (or membrane receptors) and the condensate components. While for glycinin droplets the inclusion of charges in the membrane promotes dewetting, for other condensates the presence of charges could favour wetting. This is exemplified by Dengue and Zika virus capsid proteins, which form condensates with DNA/RNA and wet negatively charged membranes^12^. To demonstrate this, we performed experiments showing wetting transitions with condensate droplets made of a polyamine and adenosine triphosphate (ATP), as well as from the oligopeptide pair poly-lysine (K_10_) and poly-aspartic acid (D_10_), see Figs. S8 and S9. All three regimes (complete and partial wetting as well as dewetting) can be observed suggesting a universal pattern. The transitions in these highly charged systems (positive for PDDA/ATP droplets^26^ and negative for K_10_/D_10_ condensates^54^) can also be modulated by membrane composition and salinity as demonstrated for glycinin, albeit at different conditions (compare Fig. 3 and Figs. S8 and S9). These results strongly indicate a widespread behavior and show that depending on the chemical nature of the condensates, different parameters should be tuned to produce wetting transitions.

## Summary and discussion

Our results demonstrate a wide spectrum of morphology changes driven by protein condensates interacting with membranes. Glycinin condensates in contact with GUVs were observed to undergo two distinct wetting transitions, with a broad intermediate regime of partial wetting (Fig. 2). We developed a theoretical framework that allows quantification of the membrane-condensate interaction and description of the system geometry by directly measuring the apparent contact angles with optical microscopy. The observed wetting behavior can be easily modulated by salinity or membrane charge, as we demonstrate for two additional condensate systems. The wetting transitions were also observed for a more complex lipid mixture containing cholesterol as a closer mimetic to biomembranes (Fig. S1c). Note that in this case the values of the geometric factor are only slightly shifted towards higher NaCl concentration (see Fig S1d) compared to those for the pure DOPC membrane shown in Fig. 2. This indicates that, for this particular condensate-membrane system, increasing the membrane complexity and including a fast-flipping molecule such as cholesterol has only a minor effect on the wetting behavior compared to increasing the net surface charge of the membrane or changing the salinity.

The fact that changes in such simple system parameters allow for fine tuning of wetting and morphology suggests that cells take advantage of local changes of cytosol and membrane composition to modulate organelle shape and interactions mediated by wetting or suppressed by dewetting. For example, the phase separation of zona occludens proteins in tight junctions depends not only on the receptor density (concentration) but also on their valency^15^. Furthermore, while epithelial junctions and focal adhesions^55^ present examples of complete wetting, processes such as the interaction of stress granules^13^ or virus protein condensates^12, 56^ with the endoplasmic reticulum or the association of stress granules to lisosomes^57^ appear to involve partial wetting, and most likely wetting transitions. All these examples emphasize the significance of our findings.

Glycinin is a major storage protein in the soybean and plays a key role in plant embryogenesis. In most cotyledons, storage proteins are accumulated in the vacuoles^58^, and recently it has been reported that they not only provide nutrients for subsequent growth but also contribute to the tonoplast (vacuolar membrane) remodeling during plant development^59^. Storage proteins form phase-separated droplets in the vacuole interior that interact with membranes presenting different wetting states, as is visible in electron microscopy images^60, 61^ (see Fig. 9a-e), and with fluorescence confocal microscopy^59^ (Fig. 9f). In the final development stage, the protein droplets undergo complete engulfment by the vacuolar membrane, forming protein bodies^60^ (Fig. 9c). Such remodeling processes are relatively slow (days) as demonstrated for protein storage vacuoles in *Arabidopsis thaliana* ^59^, presumably related to the high viscosity of storage protein droplets (also shown here for glycinin). The precise mechanisms by which these processes occur in the plant cells remain unknown, but our findings show that by tuning simple parameters such as the ionic strength or the membrane composition, wetting transitions as well as complete engulfment can be induced.

**Figure 9:**
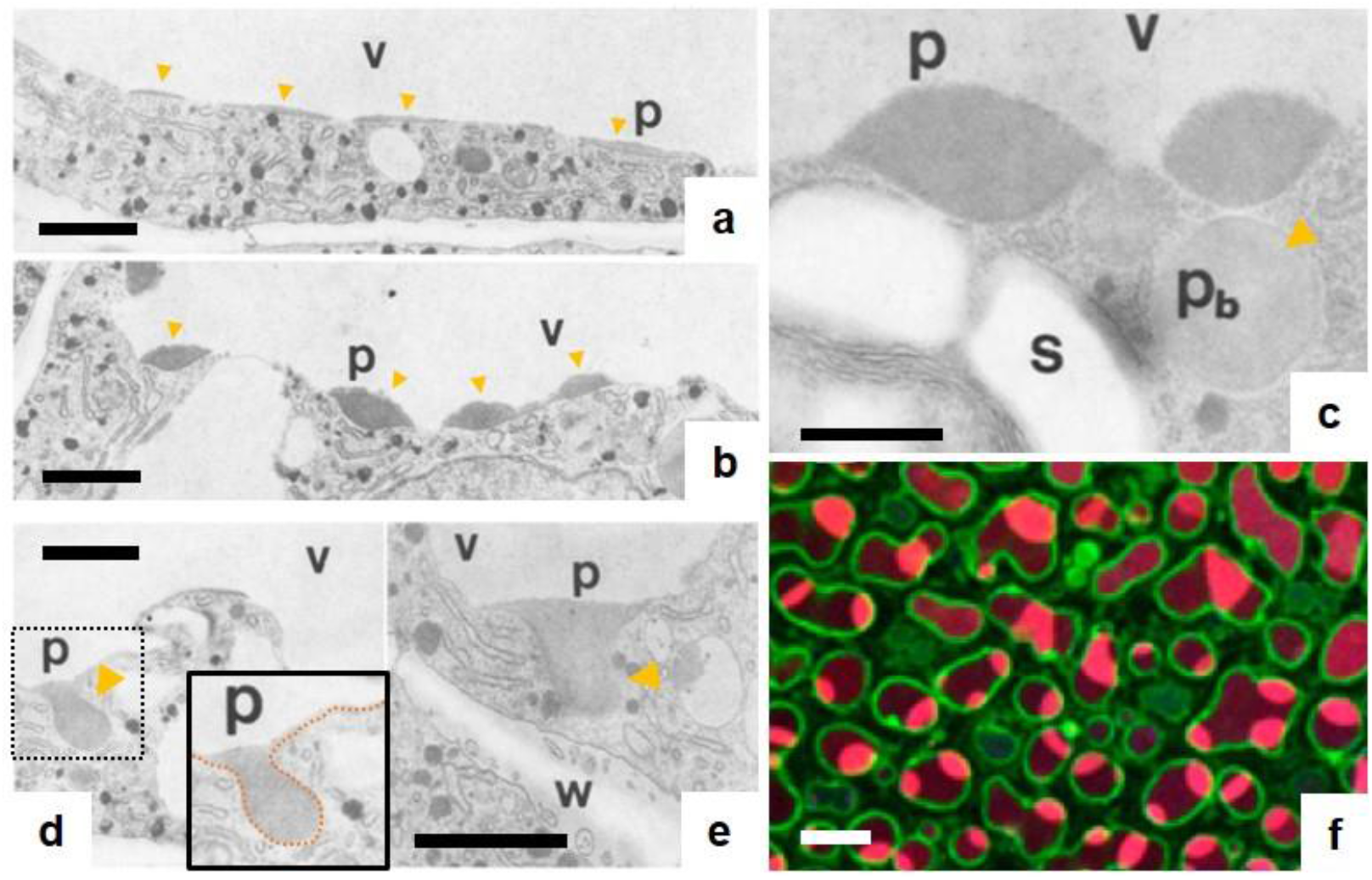
Wetting and remodeling of the tonoplast (vacuolar membrane) by storage protein condensates in plants. Electron micrographs of storage parenchyma cells of soybean cotyledons during development (a-e). Abbreviations: p=protein droplets, p_b_=protein bodies, s=starch grains, v=vacuole, w=cell wall. **(a)** Protein droplets form a thin layer on the vacuolar membrane (see darker areas to which the yellow arrows point). **(b)** Partial wetting of the protein droplets on the vacuolar membrane. Note that the membrane acquires additional curvature in the contact regions with the condensate droplets. **(c)** Protein bodies are formed as a result of complete engulfment of the protein droplets by the tonoplast and subsequent pinching off. **(d)** “Protein pockets” are formed in the vacuoles visualized as finger-like structures protruding to the cytoplasm. This is proposed to be a step prior to protein body formation_60_. The inset shows the zoomed dotted region, and the protrusion is highlighted by an orange dotted line. **(e)** Another example of protrusion formation in the vacuolar membrane. **(f)** Confocal microscopy images of protein storage vacuoles in *Arabidopsis thaliana* embryo cells showing how the protein droplets (red) wet the vacuolar membrane (green). Scale bars in (a-e) are 1 µm and in (f) 10 µm. Images (a-e) were adapted from reference ^60^ with permission from SNCSC and image (f) from reference ^59^.

Recently, wetting transitions have been reported to regulate the formation of the tight-junction belt via elongation of junctional condensates around the apical membrane interface^62^. This is yet another example that underscores the diversity of cell processes involving wetting transitions as the result of the interaction between membrane-bound and membranelles organelles.

Under certain conditions, liquid condensates can turn into hollow condensates^6, 50, 63^ that are believed to have a key role in the formation of multilayered subcellular membranelles organelles^50, 63, 64^. Here we show that membrane interaction with hollow condensates provides additional compartmentation for the system, with membrane or protein enclosed compartments. Membrane buds and necks can be formed as a result of the interplay between condensate-membrane adhesion, membrane elasticity and capillary forces (Figs. 5, S3) similarly to deformations observed in storage vacuoles^59^. Surprisingly, when excess membrane area is available, the droplet-membrane interface forms ruffles and fingers generating striking irregularly curved structures (Fig. 6), characteristic for the glycinin-vesicle system but not for the ATPS-vesicle one. The complex condensate-membrane morphologies observed for the protein condensates can be tuned by tension, or by changing the material properties of the condensates and the membrane. The diameter of the observed ruffles and fingers are around and below a micrometer (Figs. 6c,d, S5), i.e. in the dimension range of relevant mesoscale intracellular structures, like the tubular network connecting the Golgi apparatus with the endoplasmic reticulum^65^. The membrane molding observed here suggests that interactions between membrane-bound organelles and condensates can trigger a complex mutual remodeling process generating prominently curved and stable structures. In this context, it has been reported that during embryogenesis in plant cells, droplets of storage proteins, including glycinin, remodel the vacuolar membrane generating protrusions that are finger-like structures inserting in the cytoplasm^60^ as shown in Fig. 9d,e. In addition, glycinin condensation was proposed to occur in the endoplasmic reticulum for the ionic strength range used here^66^ and thus, as we demonstrate, could contribute to reshaping it. This could also be a relevant mechanism for the morphogenesis and organization of membrane-bound organelles such as the Golgi apparatus^67, 68^. Furthermore, these systems could provide templates for the design of multiphase compartments and complex connectivity to be used in synthetic biology. Plant proteins like glycinin have the advantage of being widely abundant and economical, which makes them ideal materials for the scale-up processing in biotechnological applications^6^. In this regard, synthetic protocells can now be sequentially assembled, using microfluidics and pico-injection^69^, to form droplet-stabilized GUVs and to load them with different biomolecular components^70^. Injecting biomolecular condensates into GUVs can provide a promising method to create protocells that undergo budding and form aqueous sub-compartments in a controlled and reproducible manner, allowing to host chemical or enzymatic reactions^71, 72^. By changing the interfacial tension of the condensates, the size of these compartments can be varied over several orders of magnitude, from tens of nanometers to tens of micrometers. Additionally, this system could be exploited for the development of high complexity synthetic biology systems, like the assembly of bacteriogenic protocells mediated by condensates^73^. Finally, condensate-membrane interactions triggering fingering and ruffling as shown here, demonstrate that they do not only play a role in concentration-regulated phase separation and nucleation^9^, but also dramatically mold membranes. Upon encounter with membrane-bound organelles, liquid condensates act as sculptors of intricate membrane structures, generating local curvature, without the involvement of active processes.

## Methods

### Materials

The phospholipids 1,2-dioleoyl-sn-glycero-3-phosphocholine (DOPC), Soy L-α-phosphatidylcholine (soy-PC), 1,2-dipalmitoyl-sn-glycero-3-phosphocholine (DPPC), 1,2-dioleoyl-sn-glycero-3-phospho-L-serine (DOPS), and 1,2-dioleoyl-3-trimethylammonium-propane (DOTAP), cholesterol (Chol), 1,2-dipalmitoyl-sn-glycero-3-phosphatidylethanolamine-N-(lissamine rhodamine B sulfonyl) (DPPE-Rh) as chloroform solution, galbeta1-3galnacbeta1-4(neuacalpha2-3)galbeta1-4glcbeta1-1’-cer (GM1) as powder were purchased from Avanti Polar Lipids (IL, USA). The soluble dye 2-(3-diethylamino-6-diethylazaniumylidene-xanthen-9-yl)-5-sulfo-benzenesulfonate (Sulforhodamine B) was obtained from Termofisher (MA, USA). The fluorescent lipid dye ATTO 647N-DOPE was from ATTO-TEC GmbH (Siegen, Germany). Polyvinyl alcohol (PVA, with MW 145000) was purchased from Merck (Darmstadt, Germany). Chloroform obtained from Merck (Darmstadt, Germany) was of HPLC grade (99.8 %). The lipid stocks were mixed as chloroform solutions at 4 mM, contained either 0.5 mol% ATTO 647N-DOPE (for the glycinin condensate experiments) or 0.1 mol% DPPE-Rh (for the ATPS experiments) and were stored until use at −20°C. For neutral lipid compositions in the measurements with glycinin, DOPC was employed, while for the charged membranes were prepared from DOPC:DOPS and DOPC:DOTAP mixtures. The membrane composition for the experiments with PEG-dextran ATPS was (DOPC:GM1:DPPE-Rh, 95.9:4:0.1 mol%).

Polydiallyldimethylammonium chloride (PDDA, 200-350 kDa, 20 wt% solution in H2O), adenosine triphosphate (ATP), fluorescein isothiocyanate isomer (FITC), sucrose, glucose, dimethyl sulfoxide (DMSO), Tris HCl buffer, potassium chloride (KCl), magnesium chloride (MgCl_2_), sodium hydroxide (NaOH) and sodium chloride (NaCl) were obtained from Sigma-Aldrich (Missouri, USA). Gm1PEG (average molecular weight 8000 g/mol) and dextran from Leuconostoc mesenteroides (molecular weight between 400 kDa and 500 kDa) were purchased from Sigma-Aldrich. The oligopeptides poly(l-lysine hydrochloride) (degree of polymerization n = 10) (K10) and poly(l-aspartic acid sodium salt) (degree of polymerization n = 10) (D10) were purchased from Alamanda Polymers (AL, USA) and used without further purification (purity≥95%). A N-terminal TAMRA-labelled version of K10 was purchased from Biomatik (Ontario, Canada). All solutions were prepared using ultrapure water from SG water purification system (Ultrapure Integra UV plus, SG Wasseraufbereitung) with a resistivity of 18.2 MΩ cm.

### Protein purification

Preparation of glycinin was achieved as described in Chen et al.^6^. Briefly, the defatted soy flour was dispersed in 15-fold water in weight and adjusted to pH 7.5 with 2 M NaOH. This slurry was then centrifuged (9000×g, 30 min) at 4°C. Dry sodium bisulfite (SBS) was added to the supernatant (0.98 g SBS/L), the pH of the solution was adjusted to 6.4 with 2 M HCl, and the obtained turbid dispersion was kept at 4 °C overnight. After that, the dispersion was centrifuged (6500×g, 30 min) at 4 °C. The glycinin-rich precipitate was dispersed in 5-fold water and the pH was adjusted to 7. The glycinin solution was then dialyzed against Millipore water for two days at 4 °C and then freeze-dried to acquire the final product with a purity of 97.5%_6_.

### Protein labeling

A 20 mg/mL soy glycinin solution was prepared in 0.1 M carbonate buffer (pH 9). FITC was dissolved in DMSO at 4 mg/mL. The FITC solution was slowly added into the protein solution with gentle stirring to a final concentration of 0.2 mg/mL. The sample was incubated in the dark while stirring at 23 ºC for 3 h. The excess dye was removed using a PD-10 Sephadex G-25 desalting column (GE Healthcare, IL, USA), and buffer was exchanged for ultrapure water. The pH of the labeled protein solution was adjusted to 7.4 by adding 0.1 M NaOH. For fluorescence microscopy experiments, an aliquot of this solution was added to the working glycinin solution to a final concentration of 4%.

### Glycinin condensates formation

A glycinin solution at 20 mg/mL was freshly prepared in ultrapure water and pH was adjusted to 7. The solution was filtered to remove any insoluble materials using 0.45 µm filters and was kept at 4°C before use. To form the condensates, 50 µL of glycinin solution were mixed with the same volume of a NaCl solution of twice the desired final concentration, to obtain a 10 mg/mL solution of glycinin condensates. At this protein concentration, condensates size is optimum for microscopy and distances are big enough so they do not coalesce into a large condensate during the experiment.

### Hollow condensates

Salinity shifts in a coacervate suspension towards the phase-coexistence boundary, induced either by decreasing or increasing the salt concentration, results in hollow condensate formation by triggering the formation of protein-poor phase within already formed condensates as previously reported^6^. Here, to generate hollow condensates, first 2 mL of the soy glycinin solution containing 10% (w/w) FITC-labeled protein was mixed with equal volume of 200 mM NaCl solution to induce condensate formation at 100 mM NaCl final concentration. Then, this condensates suspension (100 µL) was mixed with an equal volume of pure water to induced hollow condensates formation at 50 mM NaCl final salinity. To produce hollow condensates at 150 mM, the condensate suspension was mixed with an equal volume of 200 mM NaCl solution and then diluted with 150 mM NaCl solution once more. These steps trigger phase separation within the preformed condensates creating a protein-poor phase within them^6^.

### PDDA/ATP droplets formation

The phase separated droplets were formed by gently mixing aliquots of stocks solutions of Tris HCl (pH= 7.4), MgCl_2,_ glucose, PDDA and ATP (following this order), to a final volume of 20 µL. For labeling, a 0.5%mol of the water-soluble dye Sulforhodamine B was added. The final concentration of each component was: 20 mM Tris HCl, 5 mM MgCl_2_, 170 mM glucose, 14.8 mM ATP and 4.9 mM PDDA. The final osmolarity of the mixture was ≈ 200 mOsm.

### Oligopeptides K10/D10 droplets formation

Phase separation was triggered by gently mixing aliquots of stocks solutions of KCl, MgCl_2,_ glucose, D_10_ and K_10_ (following this sequence), to a final volume of 20 µL. For labeling a 0.1%mol of TAMRA-K_10_ was added. The final concentration of each component was: 15 mM KCl, 0.5 mM MgCl_2_, 170 mM glucose, 2mM D_10_ and 2mM K_10_. The final osmolarity of the mixture was ≈ 200 mOsm.

### Lipid membranes

GUVs were grown using the electroformation method^74^. In brief, 3-4 µL of the lipid stock were spread on two conductive indium tin oxide glasses and kept under vacuum for 1 h. The two glass electrodes were separated by a 2-mm-thick Teflon frame, forming the electroformation chamber. The lipid films were hydrated with 2 ml of a sucrose solution, matching the osmolarity of the NaCl solution in which the condensates were formed. The osmolarity was adjusted using a freezing point osmometer (Osmomat 3000, Gonotec). An electric AC field (1V, 10 Hz, sinusoidal wave) was applied for one hour at room temperature. Once formed, vesicles were diluted 1:1 in a glucose solution of the same osmolarity, and the suspension of the giant vesicles was stored at room temperature until use. The vesicles were prepared freshly before each experiment. For the experiments with PEG-dextran ATPS, the dried lipid films were hydrated in solution of PEG-rich phase as explained further below.

For the experiments in Fig. S7, the GUVs were prepared with the PVA gel-assisted method^75^ which allows vesicle swelling in high salinity conditions. Briefly, two coverslips were cleaned with water and ethanol and dried under nitrogen. PVA solution was prepared by diluting PVA in deionized water to a final concentration of 40 mg/mL. A small aliquot (20-50 µL) of the PVA solution was spread on the glass slides and dried for 1 h at 60°C. 3-4 μL of the lipid stock solution were deposited on the PVA-coated glass, and kept for 1 h under vacuum at room temperature. The chamber was assembled with a 2 mm-thick Teflon spacer, and filled with 1 ml of 150 mM NaCl solution. After 30 minutes the vesicles were harvested carefully in order to prevent PVA detaching from the cover glass.

### Condensates-membranes suspensions

A 1:10 dilution of the vesicle suspension was made in a solution of the desired final NaCl concentration. The glycinin condensate suspension of the same NaCl concentration was diluted 1:4 (with a solution of NaCl at the same concentration) and added to the vesicle suspension at a 15% v/v. After gently mixing the vesicle-condensate suspension, an aliquot was placed on a coverslip (26×56 mm, Waldemar Knittel Glasbearbeitungs GmbH, Germany) for observation with confocal microscopy. The coverslips were previously washed with ethanol and water, and passivated with a 10 mg/mL BSA solution.

For the hollow condensates interaction with vesicles, 20 µL of the hollow condensate suspension was mixed with equal volume of the 10 times diluted soy-PC vesicle solution in the same salt concentration.

For the deflation/inflation experiments, vesicles were electroformed in 200 mM, 300 mM or 450 mM sucrose and then diluted in 150 mM NaCl. Condensates were prepared at 150 mM NaCl and mixed with the vesicle dilution previously described. The vesicles were visually inspected for fluctuations and floppiness (in the absence of condensates) or interface ruffling (when in contact with glycinin droplets). At least 60 vesicles were inspected in each condition, and the experiment were performed three times.

For the interaction of membranes with PDDA/ATP or K_10_/D_10_ condensates, the vesicle suspension was diluted 1:10 into the final buffer of the corresponding droplets suspension. An aliquot of this diluted vesicle solution was then mixed with the droplet suspension in 8:1 volume ratio directly on the cover glass and sealed for immediate observation under the microscope.

### PEG-dextran ATPS and membranes

Solution preparation and mixing with GUVs followed exactly the procedure described in ^34^. Briefly, polymer solution composed of 6.87 wt % PEG and 2.86 wt % dextran (1.25 wt % of the total dextran was labeled with fluorescein isothiocyanate) were prepared and left for 2.5 days to completely phase separate. The vesicles were prepared in the PEG-rich phase and a small aliquot of the dextran-rich phase was mixed with the vesicle solution. The mixing chamber was gently shaken to break the dextran phase into small droplets. Imaging was done after 2 hours or on the next day.

### Confocal microscopy and FRAP

Confocal SP5 or SP8 microscopes equipped with a 63×, 1.2 NA water immersion objective (Leica, Mannheim, Germany) were used for imaging. FITC and ATTO 647N-DOPE were excited using the 488 nm and 633 nm laser lines, respectively. FRAP measurements were performed on the SP8 setup equipped with a FRAP booster. Circular region of interest (ROI) with diameter of 2 μm on the membrane was bleached during 3 iterative pulses of total time ∼3 s. Fluorescence intensities from ROI corresponding to photobleaching were analyzed using ImageJ. Curves were fitted using the formula:*y*=(*I*_0_+*I*_*max*_(*x*/τ_1/2_))/(1+*x*/τ_1/2_), where *I*_*max*_ is the maximal intensity and τ_1/2_ is the halftime of recovery.

### STED microscopy

An Abberior STED setup (Abberior Instruments GmbH) based on an inverted Olympus IX83 microscope (Olympus Inc., Japan) equipped with a 60×,1.2 NA water immersion objective was used to obtain the super-resolved images. The sample was excited at 640 nm and the depletion laser corresponded to a 775 nm pulsed beam. Alignment was achieved as described previously for the setup^76^. Briefly, 150 nm gold beads (Sigma-Aldrich, USA) were observed in reflection mode to overlap the center of the excitation focus with the center of the depletion focus. Corrections for mismatches between the scattering and fluorescence modes were performed using 100 nm TetraSpeck™ beads (Invitrogen, USA). To measure the resolving power of the setup, crimson beads of 26 nm diameter (FluoSpheres™, Molecular Probe) were used. A measured resolution of ∼35 nm was achieved using 80% of the STED laser power (total laser power of 1.25 W), improving by 10-fold the lateral resolution over that of the corresponding excitation laser^76^. For our experiments, 3D STED was more suitable than 2D STED, since we could eliminate the interference of out-of-focus signal coming from the curved regions of the membrane (see Fig. S5). For the images shown in Fig. 6c and S5, the pixel size is 50 nm and the pixel dwell time is 10 µs.

### Micropipette aspiration

Micropipettes were formed by pulling glass capillaries (World Precision Instruments Inc.) with a pipette puller (Sutter Instruments, Novato, CA). Pipette tips were cut using a microforge (Narishige, Tokyo, Japan) to obtain smooth tips with inner diameter between 6-10 μm. The pipette tips were coated with 2 mg/mL solution of casein (Sigma) to prevent adhesion. Latex microspheres of 6 µm diameter (Polysciences Inc., PA, USA) were used to determine the zero-pressure level. The aspiration pressure was controlled through adjustments in the height of a connected water reservoir mounted on a linear translational stage (M-531.PD; Physik Instrumente, Germany). Images were analyzed using the ImageJ software.

### Wetting geometries and contact angles

All wetting geometries involve three aqueous solutions: the interior solution *i* within the vesicle, the exterior buffer *e*, and the glycinin condensate *c*. These aqueous solutions are separated by three surface segments (Fig. 2b). The glycinin condensate *c* is separated from the exterior buffer *e* by the *ce* interface. When the interface forms a contact line with the membrane, this line partitions the membrane into two membrane segments. The membrane segment *ic* forms the contact area with the condensate whereas the other membrane segment *ie* is exposed to the exterior buffer. At the contact line, the three surface segments form three apparent contact angles *θ*_*c*_, *θ*_*i*_, and *θ*_*e*_ that add up to 360° (Fig. 2b). These three contact angles are related to the three surface tensions via the tension triangle in Fig. 2c ^40^ and to the affinity contrast *W* as defined in Eq. (9) below.

### Contact angle determination

In order to adequately measure the contact angles between the different interfaces from microscopy projections, it is necessary that in the projected image, the rotational axis of symmetry of the vesicle-droplet system lies in the image plane (see Fig. S10). Otherwise, a wrong projection will lead to misleading values of the system geometry and the contact angles. Once the 3D image of the vesicle and the droplet is acquired (and reoriented to obtain correct projection), we consider the three spherical caps of the vesicle, the droplet and the vesicle-droplet interface, and fit circles to the contours to extract the different radii *R*_*i*_ and center positions *C*_*i*_ as defined in Fig. S10b. In this manner, all contact angles can be defined by the following expressions^37^:

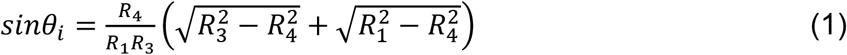

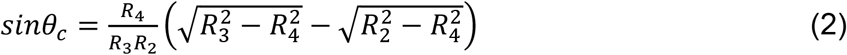

with *R*_*3*_ ≥ *R*_*2*_

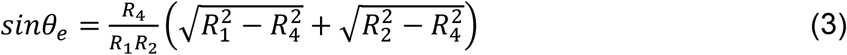

For the case of the center position *C*_*3*_ being located above *C*_1_, see Fig. S10b (and the membrane is curved towards the droplet), these relations become^37^:

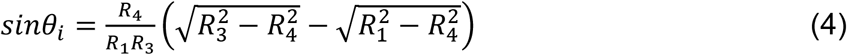

With *R*_*3*_ ≥ *R*_1_

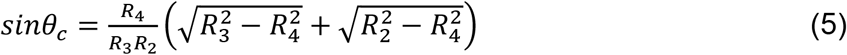

and

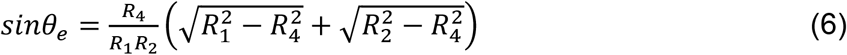

where *R*_1_, *R*_*2*_, and *R*_*3*_ are the radius of the circles fitting the vesicle, the droplet and the contact line, respectively. *R*_*4*_ is the apparent contact line radius, that can be obtained as follows:

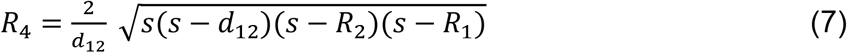

where 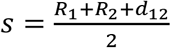. In this manner, the only input parameters required to be measured from the projection images are the radii of the fitted circles (*R*_1_, *R*_*2*_, and *R*_*3*_) and the distance between the centers of the vesicle and the droplet circles (*d*_1*2*_).

Note that the above analysis is only valid for spherical caps morphologies as shown in Figs. 2-3, and cannot be applied to ruffled membrane-droplet interfaces (Figs. 6-8), which do not fulfill the spherical cap condition.

### From contact angles to fluid-elastic parameters

The contact angles in Figs. 2-4, are related to the different surface tensions that pull the three surface segments along the contact line, see Fig. 2b,c. One of these surface tensions is provided by the interfacial tension Σ_*ce*_ of the condensate-buffer interface. The latter tension is balanced by the difference 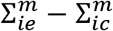 between the membrane tensions 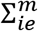 and 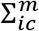 of the two membrane segments. This balance implies that the three surface tensions form the sides of a triangle (Fig. 2c). As shown in Ref^37^, the mechanical tensions of the two membrane segments are given by

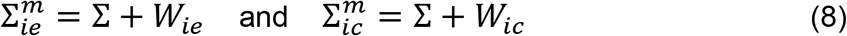

where Σ denotes the lateral stress within the membrane, which is conjugate to the total surface area *A* of the vesicle membrane, whereas *W*_*ie*_ and *W*_*ic*_ represent the adhesion free energies per unit area of the external buffer *e* and of the condensate *c* relative to the interior solution *i* ^37^. The decomposition in Eq. (8) follows from the shape functional for vesicle-droplet systems, which contains both the Lagrange multiplier term Σ*A* and the adhesion energy which depends on the adhesion parameters *W*_*ie*_ and *W*_*ic*_. The adhesion parameter *W*_*ic*_ is negative if the membrane prefers the condensate over the interior solution and positive in the opposite case. For simplicity, possible contributions from the spontaneous curvatures of the membrane segments^37^ have been ignored in Eq. (8). The affinity contrast between the condensate and the external buffer is then given by

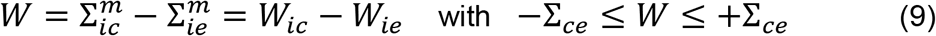

where the inequalities follow from the tension triangle in Fig. 2c and the general property that each side of a triangle must be smaller or equal to the sum of the two other sides. Note that the geometry-dependent lateral stress Σ drops out from the affinity contrast *W*, which is negative if the membrane prefers the condensate phase *c* over the external buffer *e* and positive otherwise. The limiting value *W* = −Σ_*ce*_ corresponds to complete wetting by the condensate phase whereas the limiting case *W* = +Σ_*ce*_ describes dewetting from the condensate phase (which is equivalent to complete wetting by the external buffer). The tension triangle in Fig. 2c also implies the relationships^37^

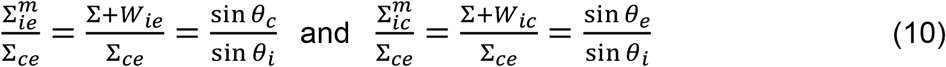

between the surface tensions and the contact angles as follows from the law of sines for the tension triangle in Fig. 2c. When we take the difference of the two equations in Eq. (10) the affinity contrast *W* in Eq. (9) becomes equal to

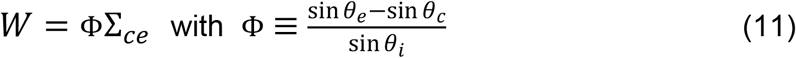

Thus, the rescaled affinity contrast *W*/Σ_*ce*_ which is a mechanical quantity related to the adhesion free energies of the membrane segments, is equal to the factor Φ, which is a purely geometric quantity that can be obtained from the three contact angles as determined by analysing the optical images (Fig. 2a,e). The inequalities in Eq. (9) imply the inequalities −1 ≤ Φ ≤ 1 for the geometric factor Φ where the interpretation for the limiting values of Φ follows from the limiting cases for the affinity contrast *W* in Eq. (9). The smallest possible value Φ = −1 corresponds to complete wetting of the membrane by the condensate phase; the largest possible value Φ = +1 to dewetting of the membrane by this phase. The dimensionless factor Φ is negative if the membrane prefers the condensate over the exterior buffer and positive otherwise. It then follows from the data in Fig. 2d that the membrane prefers the condensate for molar salt concentrations *X > X*_0_ ≃ 93 mM and the exterior buffer for *X* < *X*_0_. For *X* = *X*_0_, the geometric factor Φ vanishes, corresponding to equal contact angles *θ*_*c*_ = *θ*_*e*_ and no affinity contrast between the condensate and the exterior buffer (*W* = 0). Therefore, by measuring the geometric factor Φ as a function of the salinity or another control parameter, we obtain the affinity contrast 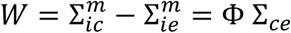 in units of the interfacial tension Σ_*ce*_ as a function of this control parameter. Note that the geometric factor is scale-invariant and does not depend on the sizes of a given vesicle and condensate pair, as exemplified in Fig. S11.

### Criterion for ruffling of the condensate-membrane interface

Ruffling and fingering of the *ic* interface implies an increase of the bending energy (assuming negligible spontaneous curvature). Ruffling will be observed when this bending energy is overcompensated by the gain in adhesion energy for transferring membrane area from the *ie* (condensate-free) segment to the *ic* interface (in contact with the condensate). When we transfer the membrane area ∆*A*, the gain in adhesion energy is *E*_*ad*_ = *W*∆*A*. Introducing Eq. (11) yields *E*_*ad*_ = ΦΣ_*ce*_∆*A*, which is negative for negative Φ; note that ruffling is observed only at for [NaCl] ≥ 100 mM where Φ < 0, see Fig. 1. The bending energy *E*_*be*_ is proportional to 8*πκ* and also proportional to the number *N*_*pro*_ of protrusions formed by the membrane area ∆*A*. The total energy change associated with the transferred membrane area is then given by *ΔE* = *E*_*ad*_ + *E*_*be*_ = ΦΣ_*ce*_∆*A* + *c8πkN*_*pro*_. Here Φ < 0 and *c* is a dimensionless coefficient of the order of 1. This energy change *ΔE* is negative for |Φ|Σ_*ce*_∆*A > c8πkN*_*pro*_ (where |Φ| is the absolute value of Φ). Ignoring the dimensionless coefficient *c*, we obtain the simple criterion

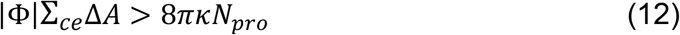

for the area transfer and ruffling to be energetically favorable. To examine this relation, we consider the vesicle condensate pair shown in Fig. 8b. For the interfacial tension and the bending rigidity we take Σ_*ce*_ *⋍*0.5 mN/m (as measured here) and *k ⋍*10^−19^ J. The area change stored in ruffles that can be pulled out with a micropipette is of the order of 50 μm^2^. For the salinity conditions where ruffling is observed ([NaCl] ≥ 100 mM), |Φ| ranges between 0.2 and 1, see Fig. 1. If we take for the number of protrusions *N*_*pro*_ =100, which is a generous overestimate, we still obtain that the criterion Eq. (12) is satisfied whereby the adhesion energy gain is still orders of magnitude higher than the bending energy penalty for ruffling.

Judging from the images in Figs. 6-8, the fingers and undulations do not have characteristic length scale. Their dimensions are defined by the available membrane area (∆*A*) to form the protrusions and the volume constraints imposed by the condensate and vesicle sizes.

### Glycinin condensates material properties

Measuring the material properties of biomolecular condensates is a challenging task, that typically relies on fluorescence recovery after photobleaching (FRAP) measurements and on quantifying the coalescence kinetics of two condensate droplets^2, 77, 78^. These techniques can provide information about the viscosity^77, 78^ and can yield the inverse capillarity number (i.e. the ratio of surface tension/viscosity)^2^, respectively. Recently, the micropipette aspiration method has been applied to quantify both, the viscosity and the surface tension of condensate droplets, provided that they behave as Newtonian fluids at the experimental time scale (>1s)^47^.

In a previous study, glycinin condensates were shown to exhibit only negligible fluorescence recovery in FRAP experiments, suggesting a highly viscous environment^6^. By using the condensate coalescence assay, the inverse capillary number of glycinin droplets was measured to be *η*/Σ _*ce*_ ≈ 9.69 s/μm ^6^. A very rough estimate of the surface tension based on the molecular size of glycinin yielded the value of Σ_*ce*_ ≈ 0.16 mN/m, implying for the viscosity η≈1.6 kPa.s ^6^. Here, we attempted to measure directly the viscosity and surface tension by means of the micropipette aspiration method. However, as can be observed in Fig. S12, glycinin condensates do not flow inside the pipette. The aspiration can only proceed until a pressure value (the maximum suction pressure achieved by our setup is up to 2500 Nm^-2^) beyond which the pipette gets clogged and the condensate cannot be further aspirated nor released. This outcome does not depend on the condensate/pipette diameter ratio. In addition, aging effects were discarded since they occur at time scales (days) ^6, 79^ longer than those required for our experiments (minutes-hours). Pipette clogging prevented us from measuring the viscosity, but we could estimate the tension by means of the Laplace equation 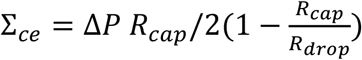, where ∆*P* is the applied pressure, and *R*_*cap*_ and *R*_*drop*_ are respectively the radius of the spherical cap formed by the part of the droplet inside the pipette and the radius of the droplet outside the pipette (see sketch in Fig. S12b). The tension value obtained in this way for glycinin droplets in the presence of 100 mM NaCl is Σ_*ce*_= (0.5 ± 0.3) mN/m (n=5), which is of the same order of magnitude of the value previously estimated from the coalescence assay. The condensate viscosity we obtain is η≈4.8 kPa.s. The glycinin surface tension value is similar to that of LAF-1^80^ and PolyR^81^ condensates, and the viscosity, close to that of the nucleolus^82^, as is summarized in Fig. 1 of the work of Wang et al.^47^ This particular combination of high surface tension and high viscosity might be related to the structural characteristics of these condensates. Whilst more coacervation-prone proteins feature mostly one type of polypeptide^83, 84^, glycinin is a hexamer^85, 86^ implying that it is characterized by a more bulky and complex structure. This could explain the reduced diffusion and high viscosity of the condensates.

## Supplementary Information

Figs. S1-S12 and movies S1-S5.

## Author contributions

A.M. performed most of the experiments. N.C. purified the protein, performed the experiments on hollow condensates, and pilot experiments to find the optimum conditions for condensate-membrane interaction. Z.Z. aided with STED imaging and provided ATPS contact angle measurements. A.M. analyzed the data. R.L developed the theoretical framework. R.D. supervised the project. A.M., R.L., and R.D. wrote the paper, with input from the rest of the authors.

## Acknowledgments

A.M. acknowledges support from Alexander von Humboldt Foundation. N.C. acknowledges funding from the National Natural Science Foundation of China (No. 32101972). Z.Z. acknowledges support from Grant SarsRapid by the Bank of Thuringia and CRC 1278 Polytarget. We acknowledge Y. Li for sharing images of vesicles in the presence of PEG-dextran ATPS. Some parts of this study were inspired by work performed in the MaxSynBio consortium, which was jointly funded by the Max Planck Society and the German Federal Ministry of Education and Research (BMBF). In particular, R.L. and R.D. are thankful for discussions and insightful comments on the manuscript by A.A. Hyman and T.M. Franzmann.

## Data availability

The authors declare that the data underlying the findings of this study are available within the paper and its Supplementary Information files.

