## Supporting information for "Wetting and complex remodeling of membranes by biomolecular condensates"


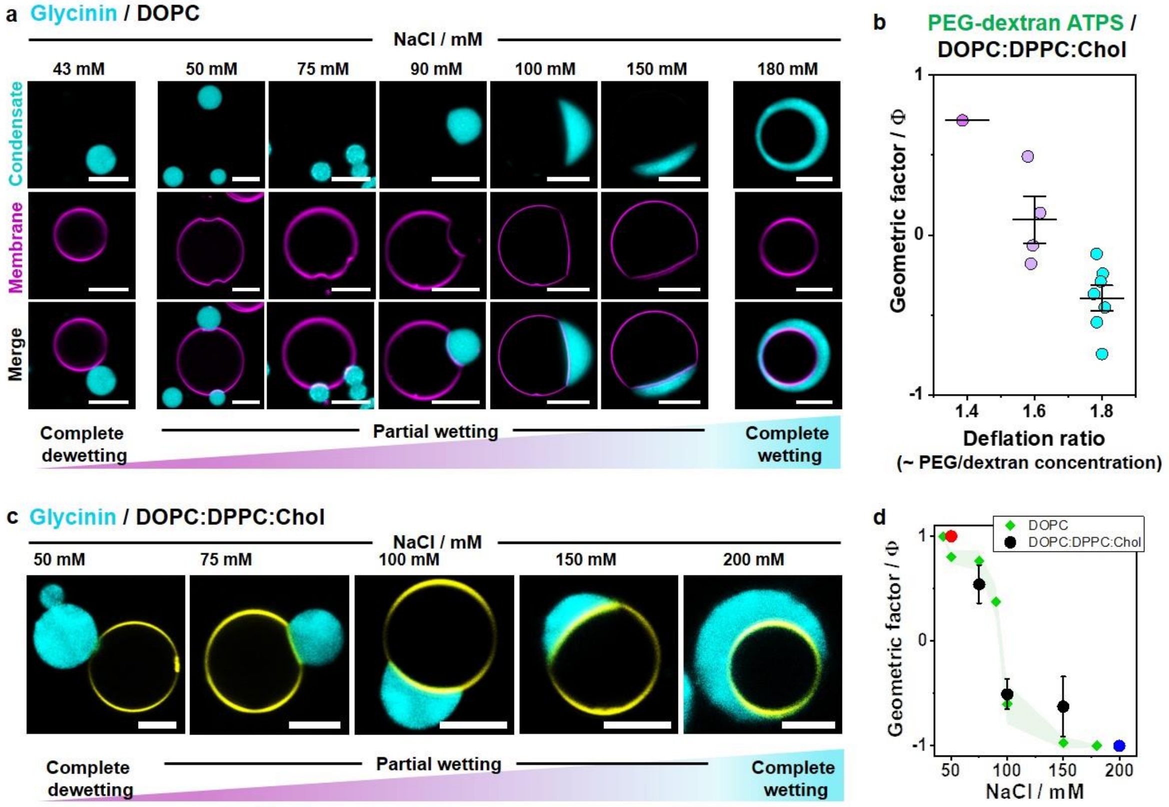


Figure S1. **Membrane wetting by glycinin condensates and PEG/dextran ATPS as a function of salinity and polymer concentration, respectively.** **(a)** Distinct wetting morphologies for glycinin condensates (cyan) in contact with giant vesicles (magenta) made of zwitterionic lipids (DOPC) at different salt concentrations. The confocal microscopy images reveal three wetting regimes separated by two wetting transitions; from complete dewetting to partial wetting (between 43 and 50 mM NaCl), and from partial to complete wetting (between 150 and 180 mM NaCl). Scale bars are 10 µm. **(b)** Geometric factor for the PEG/dextran system wetting the GUV membrane (DOPC:DPPC:Chol, 64:15:21) from the inside. The values were calculated from experimental data in Zhao et al[1]. In ATPS encapsulated in vesicles, the wetting transitions can be produced by increasing the osmolarity of the exterior medium[2]. This raises the polymers concentration inside the vesicle, promoting phase separation and enhancing the interaction of the dextran-rich phase with the membrane. The deflation ratio represents the ratio of the osmolarity outside to the initial osmolarity inside the GUVs. Values corresponding to individual vesicles are plotted (colored circles), with the black bars indicating the mean and SD. **(c)** Confocal microscopy images showing different wetting morphologies for the DOPC:DPPC:Chol GUVs (same membrane composition as in b) in contact with glycinin condensates at the indicated NaCl concentrations. Scale bars: 5 µm. **(d)** Geometric factor, Φ vs NaCl concentration for DOPC:DPPC:Chol GUVs in contact with glycinin (black, red and blue circles) and the data for pure DOPC GUVs shown in Fig. 2 (green diamonds). The green shadowed area corresponds to the SD for the DOPC data from Fig. 2d in the main text. Note that the data for the ternary mixture is shifted to slightly higher NaCl concentrations compared to that for pure DOPC. The red and blue circles correspond to the respective transitions from dewetting to partial wetting and from partial wetting to complete wetting for the DOPC:DPPC:Chol membrane. All data: mean ± SD, n=10 per composition.

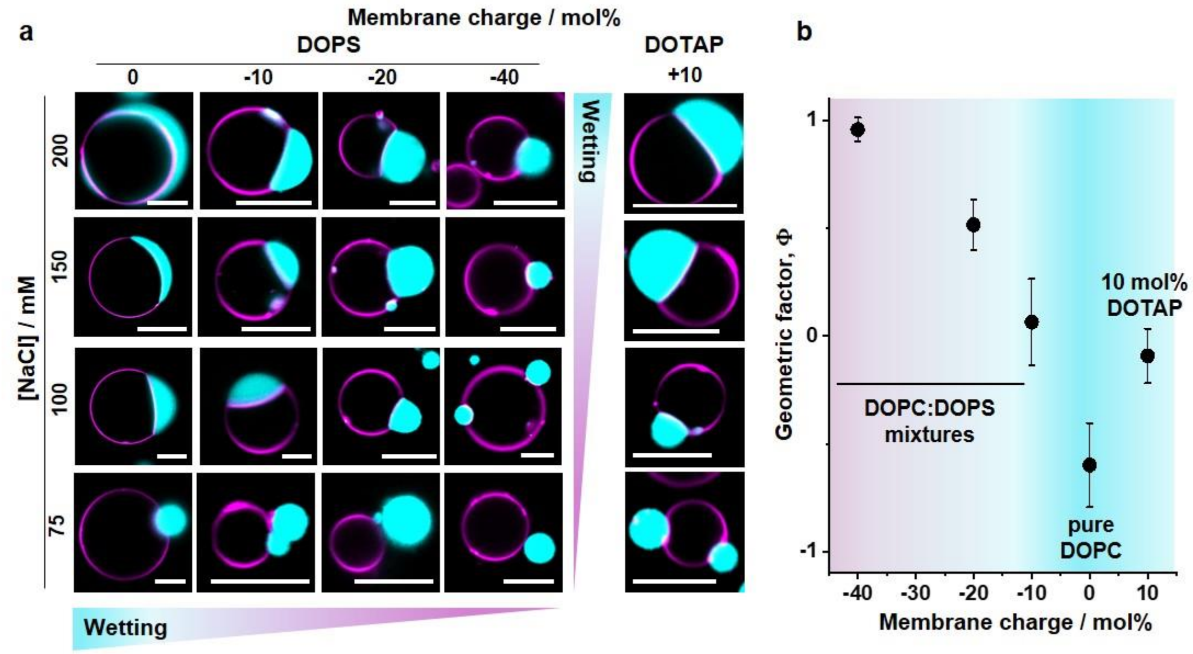


Figure S2. **Influence of membrane surface charge and salinity on the morphology of glycinin condensates (cyan) in contact with giant vesicles (magenta). (a)** Examples for wetting morphologies for DOPC vesicles containing the indicated mol % DOPS or 10 mol% DOTAP; for 200 mM and 100 mM NaCl, same data as in Fig. 3 (included here for completeness). All scale bars: 10 µm. **(b)** Geometric factor as a function of membrane surface charge at 100 mM NaCl. Negative values of the membrane charge correspond to mole fractions of DOPS in the membrane while positive ones correspond to DOTAP. Systems with 10 mol% DOPS and 10 mol% DOTAP exhibit similar morphologies and maximum wetting is observed for neutral membranes. Higher fractions of DOTAP were not explored because of the poor vesicle yield and quality for these compositions.

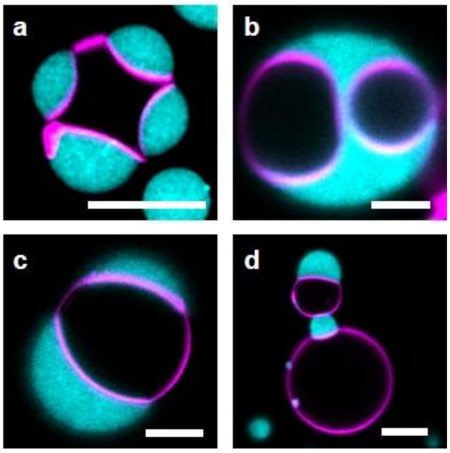


Figure S3. **Membrane molding, vesicle bridging, and compartmentation by glycinin condensates**: **(a)** Soy-PC GUV at 50mM NaCl bridging four condensates. **(b)** A single condensate engulfing two GUVs by complete wetting at 200 mM NaCl. **(c,d)** A single GUV can be wetted by more than one condensate and one condensate can bridge multiple vesicles. Scale bars: 10 µm.

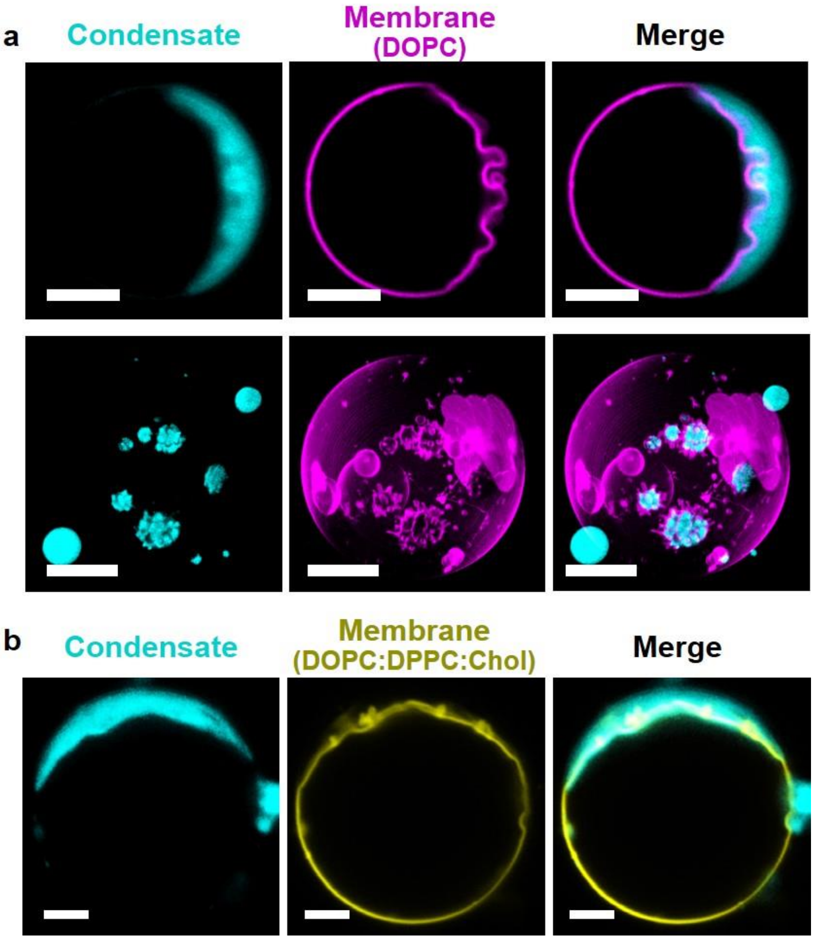


Figure S4. **Confocal cross-section and 3D reconstruction of example GUVs in contact with glycinin condensates showing different ruffling morphologies exemplified for (a) DOPC and (b) DOPC:DPPC:Chol (64:15:21) membranes.** The protrusions can point both towards the vesicle interior (as also seen in Fig. 6c) or towards the condensate, presumably depending on the volume of the droplet or vesicle provided for the buckling. The dimensions and amount of the protrusions depend on the area of excess membrane available for folding. Scale bars: 5 µm.

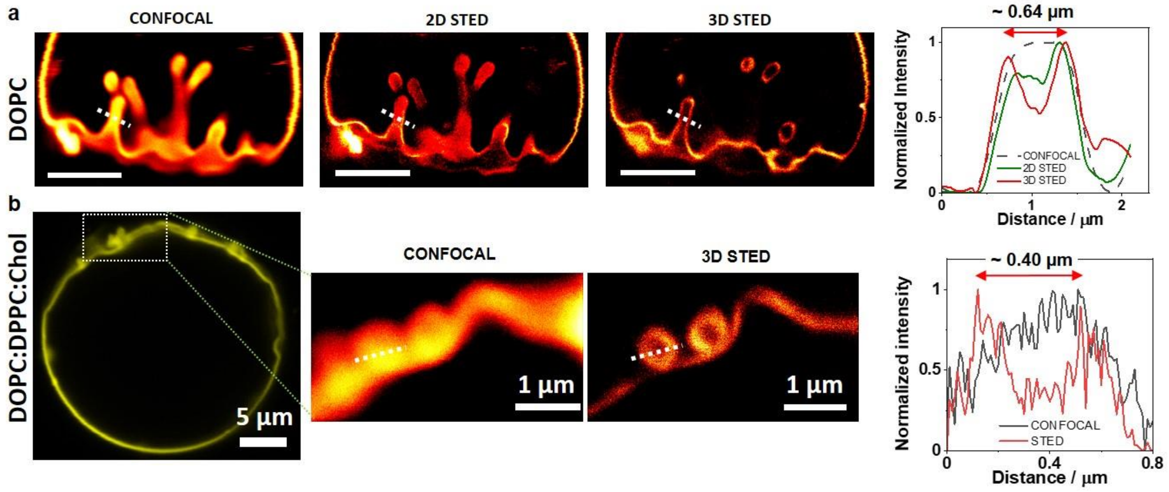


Figure S5. **3D STED helps to resolve membrane fingering eliminating interfering out-of-focus signal.** Due to the existence of protrusions and curved structures in different planes at the membrane-condensate interface, there is an increased out-of-focus signal interfering with confocal and 2D STED imaging. By using 3D STED, this interference is reduced resulting in improved z-resolution, and the membrane ruffles can be resolved. Examples of 2D and 3D STED measurements in reticulated (a) DOPC and (b) DOPC:DPPC:Chol (64:15:21) membranes. The graphs on the right show the improved resolution of 3D STED compared to 2D STED and conventional confocal microscopy, allowing the measurement of these finger-like tubes walls; the thickness of these particular finger sections is ~0.64 µm and ~0.40 µm. Scale bars in (a): 5 µm.

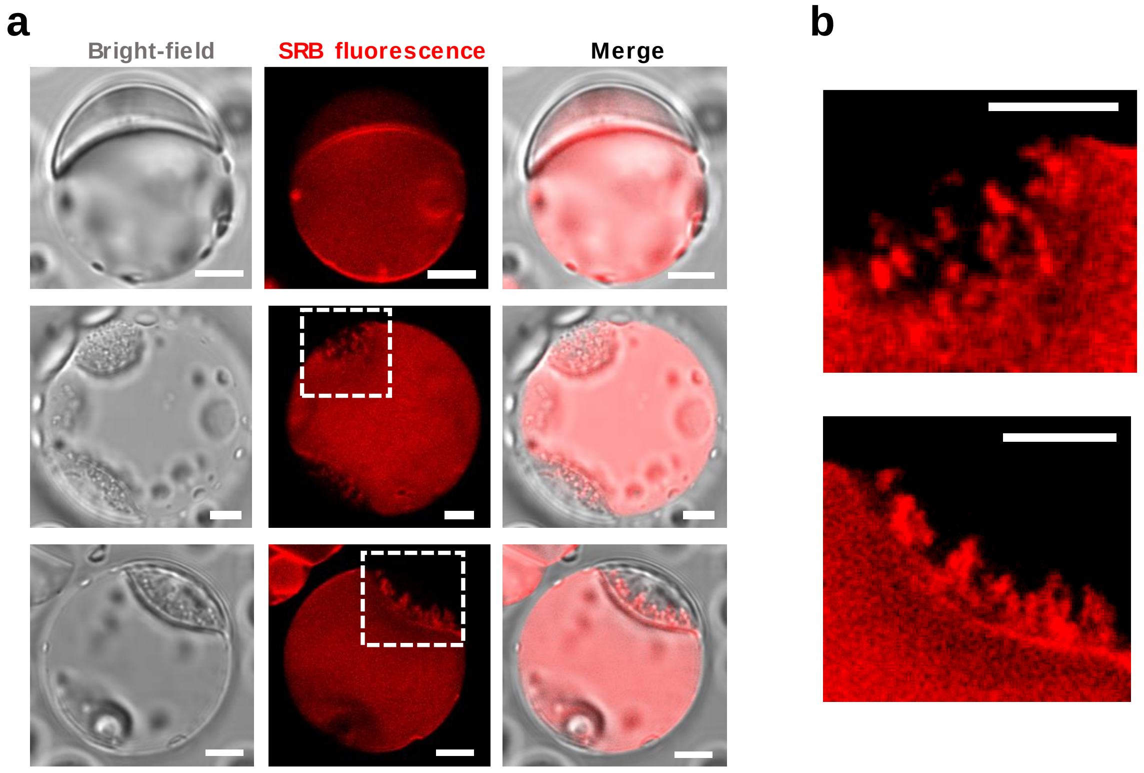


Figure S6. **Membrane integrity is preserved during wetting and ruffling.** DOPC GUVs were prepared in a solution containing the soluble dye Sulforhodamine B (SRB), in order to probe whether membrane integrity is preserved and no pores form upon condensate wetting and during the ruffling/fingering processes. **(a)** Vesicles filled with SRB do not rupture or leak during wetting and ruffling; the membrane remains intact for hours. **(b)** Zoomed images of the regions indicated with dashed lines in (a) show that the access of SRB to the finger interior is not blocked indicating that these are tube-like structures with open necks. All scale bars: 5 µm.

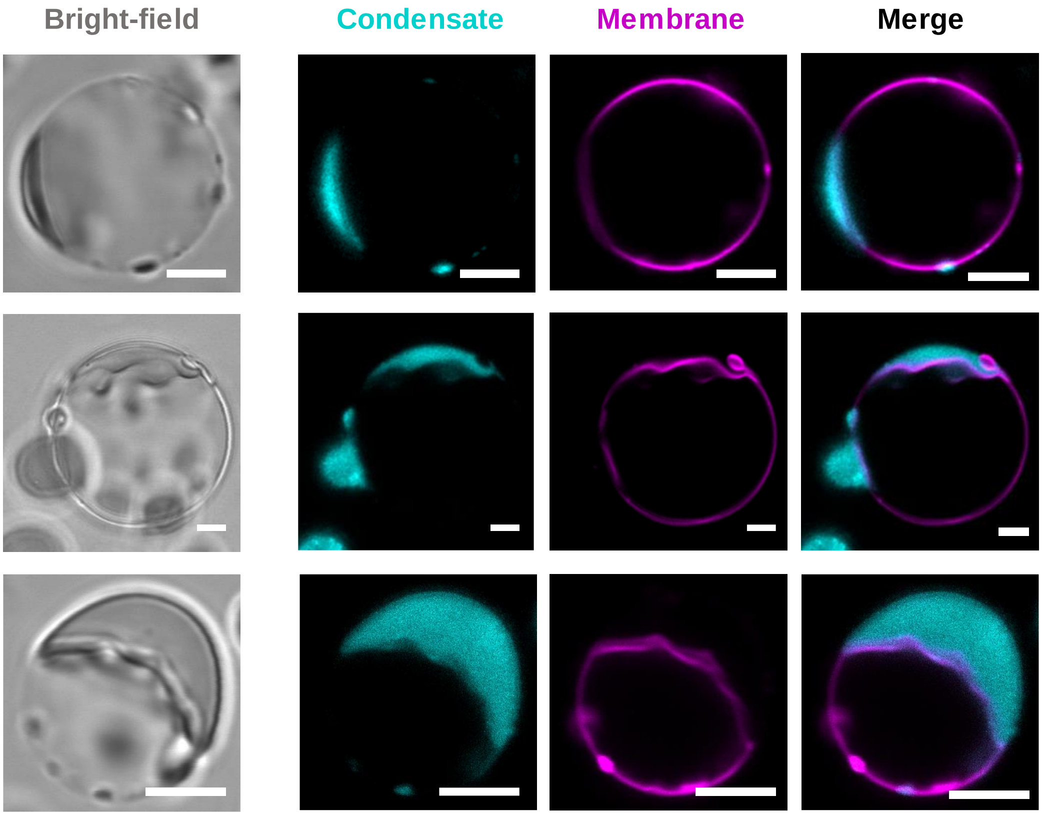


Figure S7. **Wetting and ruffling are observed in symmetric ionic conditions across the membrane.** To facilitate harvesting and microscopy observation, vesicles were typically (in all other figures) filled with a sucrose solution and diluted first in glucose and finally in NaCl, matching the osmolarity and buffer composition of the condensate suspension (see Methods). To prove that this solution asymmetry across the vesicle membrane (NaCl and glucose in the exterior vs salt-free sucrose in the interior) is not the cause of the observed ruffling as shown in other systems[3], we prepared GUVs directly in the final NaCl concentration to preserve the salinity symmetry across the membrane; vesicles were prepared using the PVA method (see Methods). In such vesicles with the same solution in the exterior and interior (150 mM NaCl) wetting and ruffling are still observed, discarding the possibility of ruffling/fingering being triggered by salt asymmetry across the membrane. Scale bars: 5 µm.

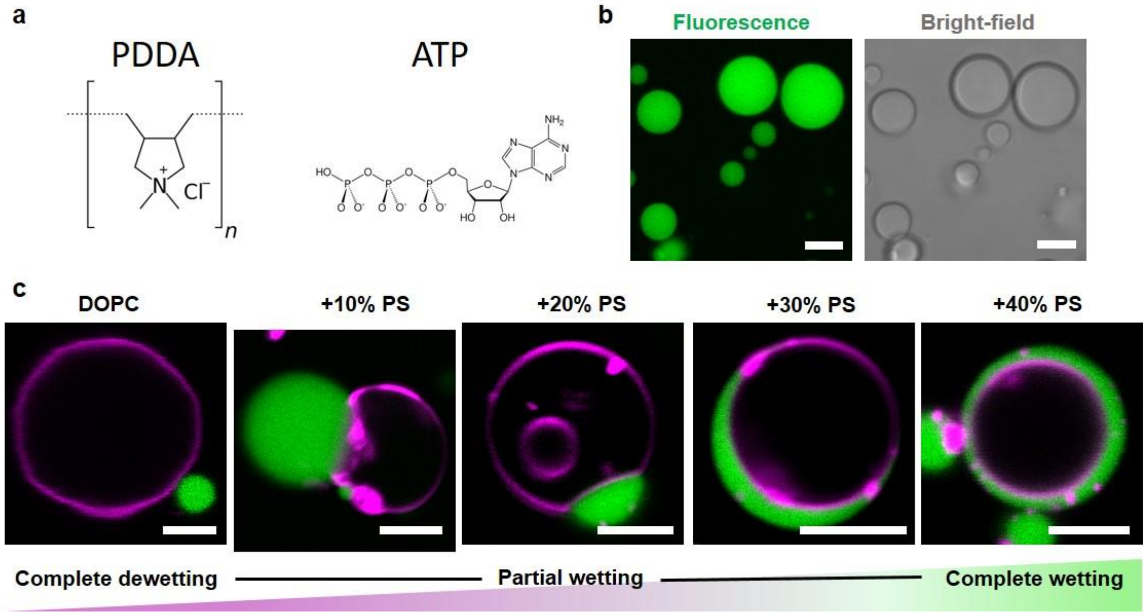


Figure S8. **Wetting transitions for pure DOPC or DOPC:DOPS GUVs in contact with PDDA/ATP droplets. (a)** Chemical structures of poly-diallyldimethylammonium chloride (PDDA, 200-350 kDa), and adenosine triphosphate (ATP). **(b)** Examples of confocal fluorescence and bright-field images of PDDA/ATP coacervates labeled with Sulforhodamine B. **(c)** Confocal microscopy images of PDDA/ATP droplets (green) in contact with DOPC vesicles (magenta) containing different molar fraction DOPS as indicated above the images. Two wetting transitions are observed for this system when changing the percentage of anionic lipids in the membrane, whereby complete wetting occurs at high fraction of the charged lipid, contrary to the behavior observed with glycinin (compare to Fig. 3a). Scale bars: 5 µm.

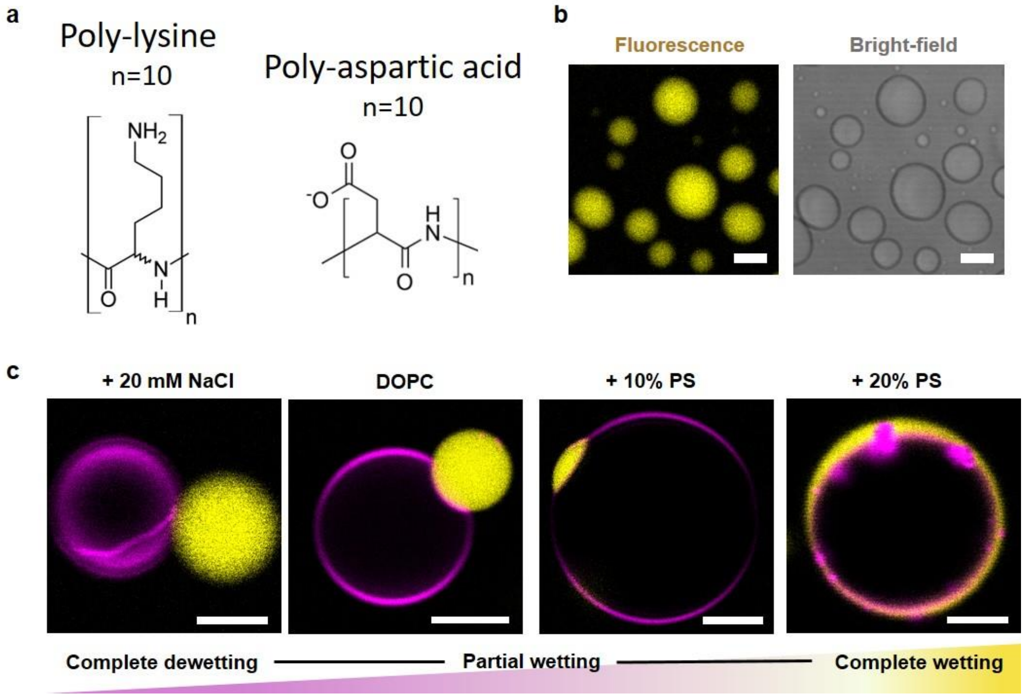


Figure S9. **Wetting transitions for pure DOPC or DOPC:DOPS GUVs in contact with poly-lysine ($K_{10}$) and poly-aspartic acid ($D_{10}$) droplets. (a)** Chemical structures of the oligopeptides poly-lysine ($K_{10}$) and poly-aspartic acid ($D_{10}$). Each peptide contains 10 monomeric units (n=10, as indicated). **(b)** Examples of confocal fluorescence and bright-field images of $K_{10}/D_{10}$ condensates labeled with TAMRA-$K_{10}$. **(c)** Confocal microscopy images of vesicles (magenta) in contact with $K_{10}/D_{10}$ droplets (yellow). The first two images are for pure DOPC vesicles in the presence of 20 mM NaCl (left) or 0 mM

NaCl (right), and the last two images correspond to a binary mixture of DOPC with the indicated % mol of DOPS in the absence of NaCl. To visualize two wetting transitions in this system it is necessary to change two parameters, namely the membrane charge to achieve complete wetting, and the salinity of the medium to achieve complete dewetting. Scale bars: 5 µm.

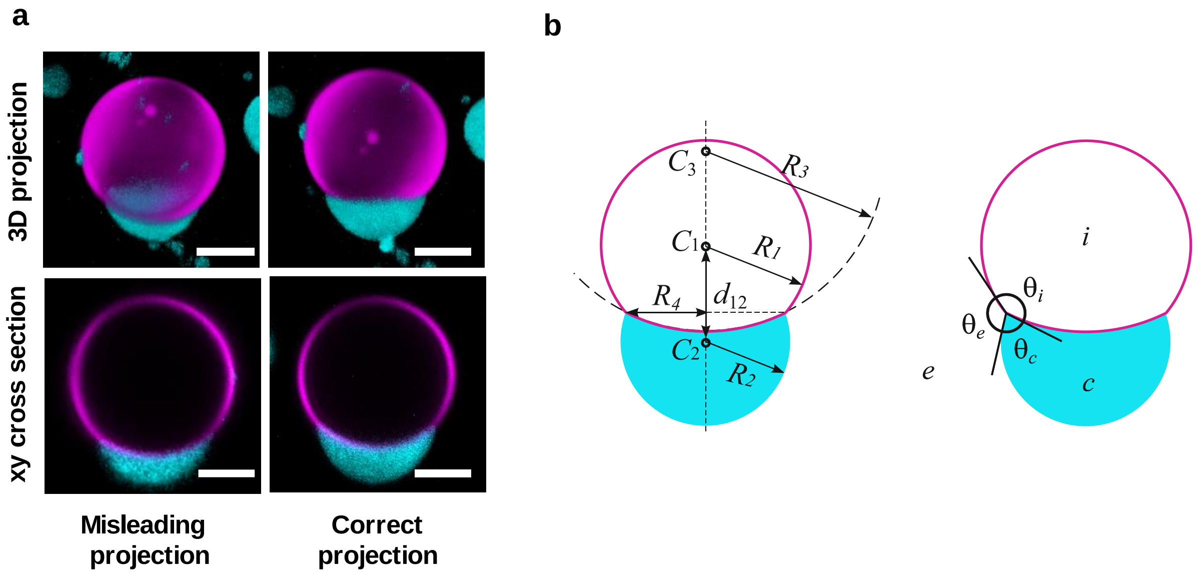


Figure S10. **Vesicle-droplet projection and parameters for measuring contact angles.** Depending on the orientation of the sample, it is very likely that the rotational symmetry axis of a vesicle-condensate couple is not parallel to the xy imaging planes, thus resulting in skewed projections in which the direct contact angle measurements would lead to misleading values. For this reason, it is necessary to correctly orient the 3D vesicle droplet confocal scan stack in order to produce the proper projection of the contact region[6]. **(a)** Examples of differently oriented 3D projections with respect to the rotational symmetry axis for the same vesicle-condensate system. On the left, the axis of symmetry passing through the centers of condensate and vesicle does not lie in the central xy cross section of the projection yielding incorrect determination of the spherical caps and contact angles, as explained in Methods. On the right, the acquired 3D projection is rotated so that the symmetry axis lies parallel to the xy cross sections. Such images are used to then obtain the geometric descriptors of the system. Scale bar: 10 µm. **(b)** Parameters needed to determine the contact angles in the system. Considering three spherical caps, and fitting circular segments to the shape contours of the droplet and of the two membrane segments, allow the determination of the radii $R_1$, $R_2$, and $R_3$, together with the positions of the centers $C_1$, $C_2$ and $C_3$, respectively. Using $R_1$, $R_2$ and $d_{12}= C_1 - C_2$, $R_4$ and the contact angles can be determined as indicated in Methods.

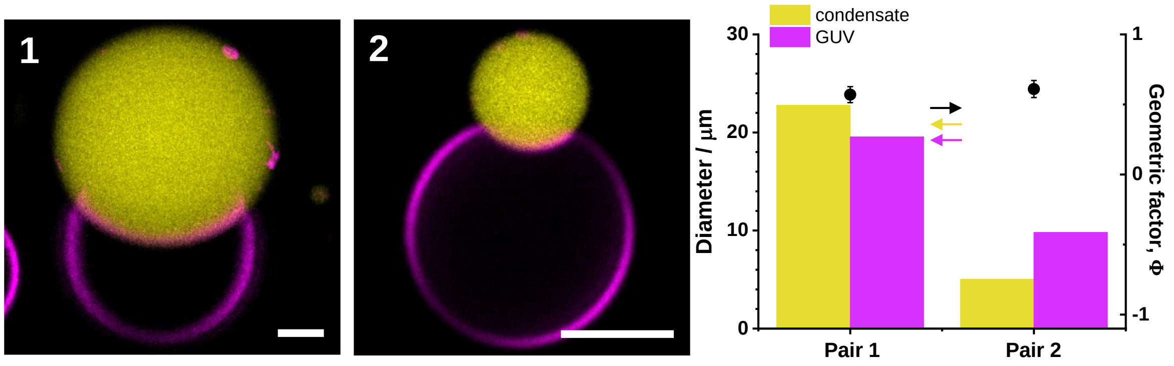


Figure S11. **Changes in the vesicle and condensate sizes do not alter the geometric factor calculation.** The interfacial tensions force balance shown in Fig. 2c is not altered by the respective sizes of the GUV and condensate interacting. The images show two examples of DOPC vesicles in contact with $K_{10}/D_{10}$ droplets. In image 1, the droplet is bigger than the vesicle and the opposite case is shown in image 2. The respective sizes of the droplets and vesicles are plotted in the graph. Despite the size differences, the force balance does not change, and the angles formed between the droplet and the membrane are the same, giving in both cases a geometric factor of Φ ≈ 0.6. Scale bars: 5 µm.

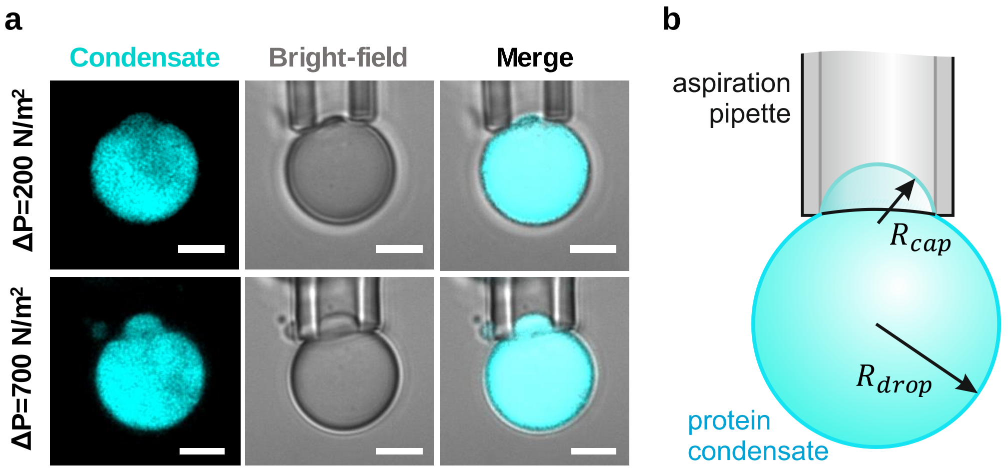


Figure S12. **Micropipette aspiration of a glycinin condensate used to assess the condensate interfacial tension $\Sigma_{ce}$. (a)** Only a small part of the condensate can be aspirated. Beyond certain pressure, the condensate cannot be further aspirated or released. Scale bar: 5 µm. **(b)** A sketch of the system with the relevant geometrical characteristics used to deduce the interfacial tension (see Methods section on glycinin condensates material properties).

### Movie Legends

**Movie S1.** Wetting dynamics for DOPC GUVs (magenta) in contact with glycinin condensates (cyan) at 100 mM NaCl (see Fig. 4a,b). Timestamps are given in minutes (hh:mm) where zero time corresponds to the initial contact between the droplet and the condensate after mixing them. The field of view is 32.6 µm × 25.9 µm.

**Movie S2.** Z-scan of a GUV-droplet system (DOPC GUV, glycinin at 100 mM) presenting ruffling/fingering of the condensate-membrane interface (see Fig. 6). Field of view is 20.0 µm × 20.0 µm.

**Movie S3.** Ruffling/fingering dynamics (DOPC GUV, glycinin at 100 mM); see Fig. 7a,b. Timestamps on the images are given in hours:minutes (hh:mm). The field of view is 26.9 µm × 26.9 µm.

**Movie S4.** Ruffling/fingering structures do not fluctuate (DOPC GUV, glycinin at 100 mM). Time is given in hours:minutes:seconds (hh:mm:ss). The field of view is 18.9 µm × 18.9 µm.

**Movie S5.** Ruffling/fingering suppression by increasing membrane tension. Micropipette aspiration of a GUV-droplet system (DOPC GUV, glycinin at 100 mM) presenting interfacial ruffling (see Fig. 8). Field of view is 28.9 µm × 28.9 µm.
